# METTL17 is an Fe-S cluster checkpoint for mitochondrial translation

**DOI:** 10.1101/2022.11.24.517765

**Authors:** Tslil Ast, Yuzuru Itoh, Shayan Sadre, Jason G. McCoy, Gil Namkoong, Ivan Chicherin, Pallavi R. Joshi, Piotr Kamenski, Daniel L. M. Suess, Alexey Amunts, Vamsi K. Mootha

## Abstract

Friedreich’s ataxia (FA) is the most common monogenic mitochondrial disease. FA is caused by a depletion of the mitochondrial protein frataxin (FXN), an iron-sulfur (Fe-S) cluster biogenesis factor. To better understand the cellular consequences of FA, we performed quantitative proteome profiling of human cells depleted for FXN. Nearly every known Fe-S cluster-containing protein was depleted in the absence of FXN, indicating that as a rule, cluster binding confers stability to Fe-S proteins. Proteomic and genetic interaction mapping identified impaired mitochondrial translation downstream of FXN loss, and specifically highlighted the methyltransferase-like protein METTL17 as a candidate effector. Using comparative sequence analysis, mutagenesis, biochemistry and cryogenic electron microscopy we show that METTL17 binds to the mitoribosomal small subunit during late assembly and harbors a previously unrecognized [Fe_4_S_4_]^2+^ cluster required for its stability on the mitoribosome. Notably, METTL17 overexpression rescued the mitochondrial translation and bioenergetic defects, but not the cellular growth, of FXN null cells. Our data suggest that METTL17 serves as an Fe-S cluster checkpoint: promoting the translation and assembly of Fe-S cluster rich OXPHOS proteins only when Fe-S cluster levels are replete.

## Introduction

Friedreich’s ataxia (FA) is a progressive neurological disorder impacting 1 in 50,000 people (Keita et al., 2022; Koeppen, 2011; Pandolfo, 2012). While the primary feature of FA is ataxia, this disease is in fact multisystemic. Patients can also develop diabetes, scoliosis, hearing and vision loss, as well as cardiomyopathy, the latter being a leading cause of premature mortality at a median age of 37.5 years (Harding, 1981; Tsou et al., 2011). FA is caused by a recessive depletion of a nuclear-encoded mitochondrial protein frataxin (FXN) (Campuzano et al., 1996), that functions as an allosteric activator of iron-sulfur (Fe-S) cluster biosynthesis (Maio et al., 2020; Parent et al., 2015; Patra and Barondeau, 2019; Srour et al., 2020). Tissue and cell samples from FA patients contain 5-30% residual FXN levels (Campuzano et al., 1997; Deutsch et al., 2010). This depletion is most often due to the expansion of a naturally occurring GAA track found within the first intron of the gene (Campuzano *et al*., 1996; Durr et al., 1996; Filla et al., 1996; Reetz et al., 2015) which triggers loss of FXN expression (Greene et al., 2007; Groh et al., 2014; Soragni et al., 2008). FA is the most common monogenic mitochondrial disease and the most common inherited ataxia. Yet we still lack approved therapies for FA, and the standard of care focuses on symptomatic management.

Fe-S clusters are ancient and universal redox cofactors (Beinert et al., 1997; Boyd et al., 2014; Braymer et al., 2021; Tsaousis, 2019). In humans, there are ∼60 known Fe-S cluster binding apoproteins (Andreini et al., 2016; Lill and Freibert, 2020) that operate throughout the mitochondrion, cytosol and nucleus. For all these subcellular compartments, cluster biosynthesis is initiated in the mitochondria where FXN accelerates Fe-S cluster formation (Parent *et al*., 2015; Patra and Barondeau, 2019; Srour *et al*., 2020). These versatile cofactors can take part in a variety of functions; while the most widely appreciated one is electron transfer, they also participate in enzyme catalysis, sulfur mobilization, and redox sensing (Braymer *et al*., 2021; Lill and Freibert, 2020; Maio and Rouault, 2020). In addition, there is evidence that in some cases Fe-S clusters contribute to the structural stability of the apoproteins into which they are integrated. Fe-S cluster binding apoproteins function in diverse cellular processes such as DNA replication and repair, nucleotide biosynthesis and energy metabolism (Maio and Rouault, 2020; Rouault, 2015). Indeed, in FA it has been well documented in patient samples as well as several disease models that there is a deficit in mitochondrial oxidative phosphorylation (OXPHOS) leading to bioenergetic defects (Gonzalez-Cabo et al., 2005; Lin et al., 2017; Lodi et al., 1999; Puccio et al., 2001; Rotig et al., 1997). It has been suggested that one reason for the neuronal and cardiac deficits observed in FA might be the high-energy demand of these tissues (Burk, 2017; Cooper and Schapira, 2003; Lynch et al., 2012).

To systematically understand the cellular consequences of FXN loss, we performed both proteomic and genetic interaction mapping on the background of FXN deficiency. Unexpectedly, we find that a consequence of FXN loss is a significant impairment in mitochondrial protein synthesis. We determine that METTL17 – a conserved mitoribosome assembly factor (Shi et al., 2019) – harbors a previously unrecognized Fe-S cluster that stabilizes its binding to the mitoribosomal small subunit. As a result, in FXN deficient cells, METTL17 activity is diminished. The data suggest that METTL17 loss is a mitochondrial repercussion of FXN deficiency and contributes to an intra-organelle translational defect.

## Results

### Loss of FXN leads to a widespread depletion of Fe-S cluster containing proteins

We sought to determine the cellular consequences of loss of FXN, an allosteric activator of Fe-S cluster biosynthesis found in the mitochondria. We performed quantitative, whole-proteome profiling of human K562 cells following CRISPR-based disruption of *FXN* versus *OR11A1,* a non-expressed gene that serves as an editing control (Fig 1A). In this analysis (Table S2), FXN itself scored as the 2^nd^ most depleted protein (Fig 1B), while its partner ISC machinery (ISCU, NFS1, LYRM4) remained unchanged (Fig S1A). On the upgoing side, IRP2, an iron regulatory protein known to be stabilized in FXN deficiency, was among the top 100 upregulated proteins. In addition, one of the top upgoing proteins was an apolipoprotein B receptor, APOBR, consistent with recent work showing a striking change in cellular lipid metabolism following the loss of Fe-S cluster biosynthesis (Chen et al., 2016; Crooks et al., 2018; Turchi et al., 2020; Wang et al., 2022). What immediately caught our attention was that of the 34 known Fe-S cluster containing proteins detected in our proteomics analysis, 33 were downregulated (Fig 1B). The only unaffected Fe-S cluster containing protein was NARF, a nuclear protein that is a component of a prelamin-A endoprotease complex (Barton and Worman, 1999). This analysis revealed that depletion of FXN leads to a near universal diminution in Fe-S cluster binding proteins. It is likely that this pervasive depletion takes place at the post-transcriptional level, as these proteins function in and are regulated by distinct pathways. The ubiquity of this depletion suggests that proper incorporation of an Fe-S cluster is essential for the structural integrity of virtually all human Fe-S cluster apoproteins.

**Fig 1:**
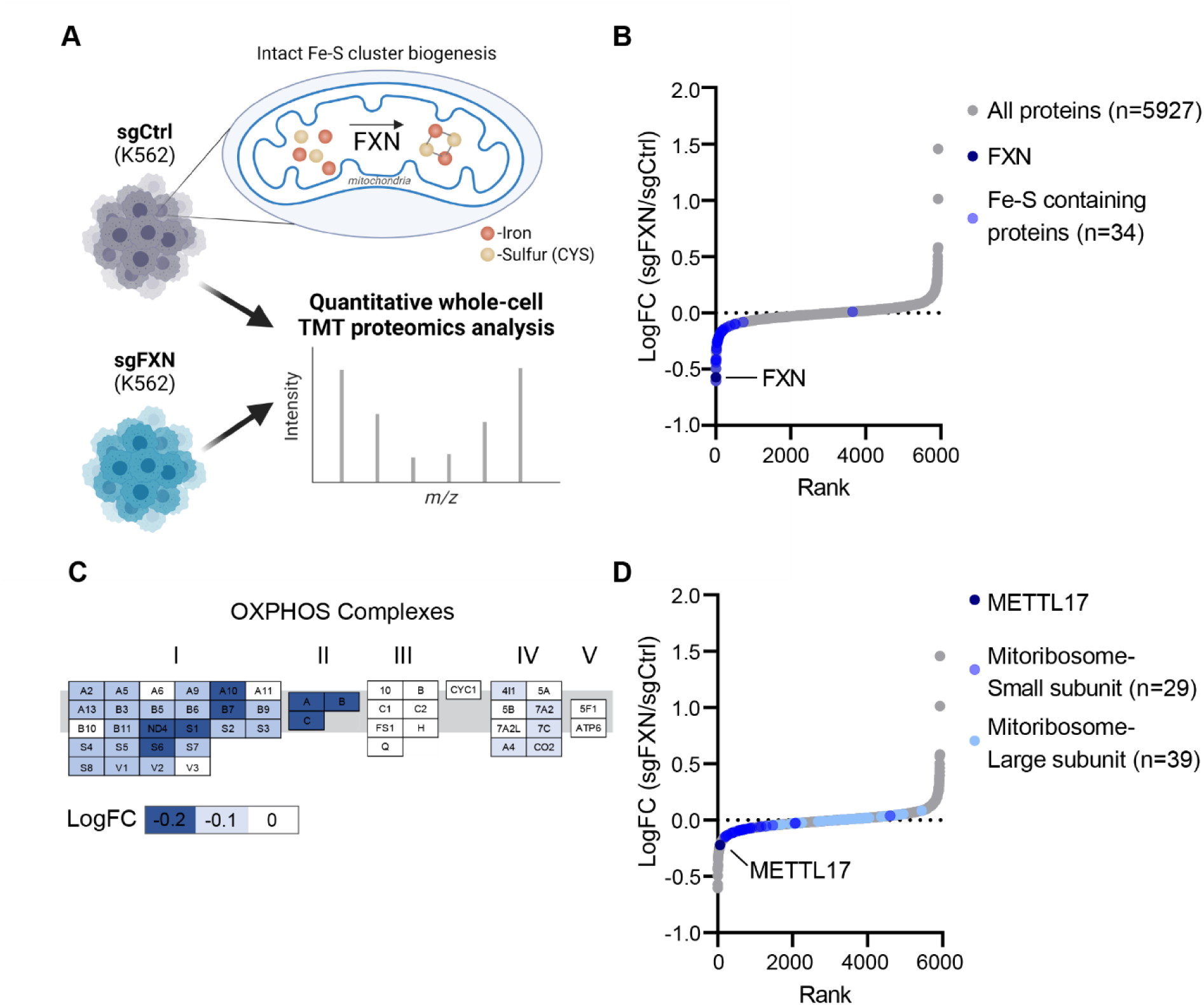
Proteomic analysis of FXN null cells reveals a marked depletion of known Fe-S cluster containing proteins and reduction of small mitoribosome subunits. A. Quantitative whole cell proteomic analysis was carried out on K562 cells edited with control or FXN targeted guides, depleting for this allosteric regulator of Fe-S cluster biosynthesis. B. Waterfall plot of protein fold change in FXN/Control cells, highlighting FXN and validated human Fe-S cluster containing proteins. C. OXPHOS proteins are organized by complex with blue indicating proteins that are depleted in FXN null cells. Genes are ordered alphabetically within complex using complex-specific prefixes (NDUF, SDH, UQCR, COX, ATP5) (e.g. A2 in CI refers to NDUFA2 whereas A in CII refers to SDHA). D. Waterfall plot of protein fold change in FXN/Control cells, highlighting proteins in the small and large mitoribosome subunit, as well as the small subunit assembly factor, METTL17.

### FXN loss results in a decrease of OXPHOS subunits as well as translational regulator METTL17

We next asked at an organelle-wide level what pathways are most altered following FXN depletion. We systematically considered all 149 MitoPathways from the MitoCarta 3.0 inventory (Rath et al., 2021) and found that in addition to the Fe-S containing proteins, the levels of respiratory chain Complex I and Complex II were significantly reduced (Fig 1C and Fig S1B). Indeed, these are the respiratory chain complexes that contain Fe-S clusters, and moreover, biopsy material from FA models will often exhibit moderate defects in these two complexes (Puccio *et al*., 2001; Rotig *et al*., 1997). Of note, our analysis did not reveal a widespread decrease in all mitochondrial-localized proteins (Fig S1C), indicating that the overall mitochondrial content is preserved despite loss of FXN.

We were intrigued to see that cells edited for *FXN* also displayed reduced levels of the mitochondrial ribosome, specifically proteins that make up the small (SSU), but not large (LSU) subunit of the mitochondrial ribosome (Fig 1D and Fig S1B). The mitoribosome is essential for the intra-organellar translation of the 13 mtDNA encoded OXPHOS subunits in humans (Christian and Spremulli, 2012; Mai et al., 2017; Ott et al., 2016), without which mitochondrial respiration cannot occur. Further examination of other proteins involved in mtDNA expression, i.e., mtDNA maintenance and mtRNA metabolism, revealed no such similar reduction in *FXN* null cells (Fig S1D-E). Intriguingly, one of the top depleted proteins in the FXN-null cells was METTL17 (Fig 1D), a seven-beta-strand methyltransferase which has recently been suggested to function as a SSU assembly factor (Shi *et al*., 2019) but has no known connections to Fe-S cluster biosynthesis. These findings indicated a profound, yet poorly fleshed out, connection between Fe-S cluster levels and mitochondrial translation which we set out to further explore.

### mtRNA translation is attenuated in the absence of FXN

We next directly tested whether FXN depletion results in a defect in intra-mitochondrial translation. To this end, we assessed the rates of protein synthesis in the mitochondria in K562 cells edited for *FXN*, *NDUFS1* or *FBXL5* (Fig 2A and Fig S2A). The latter two genes would indicate whether any changes we observe in *FXN* null cells are secondary to OXPHOS deficiency or activation of the iron-starvation response, respectively, two known consequences of FXN loss. While *FXN* deficient cells had dampened intra-mitochondrial protein synthesis, we did not observe a comparable effect in *NDUFS1* or *FBXL5* deficient cells. Furthermore, *FXN* null cells were comparatively resistant to chloramphenicol (Fig S2B), which directly inhibits mitochondrial translation. This translation defect was not a secondary consequence of a defect in mtDNA homeostasis or expression, as mtDNA copy number and mtRNA levels were intact upon loss of FXN (Fig S2C-D).

**Fig 2:**
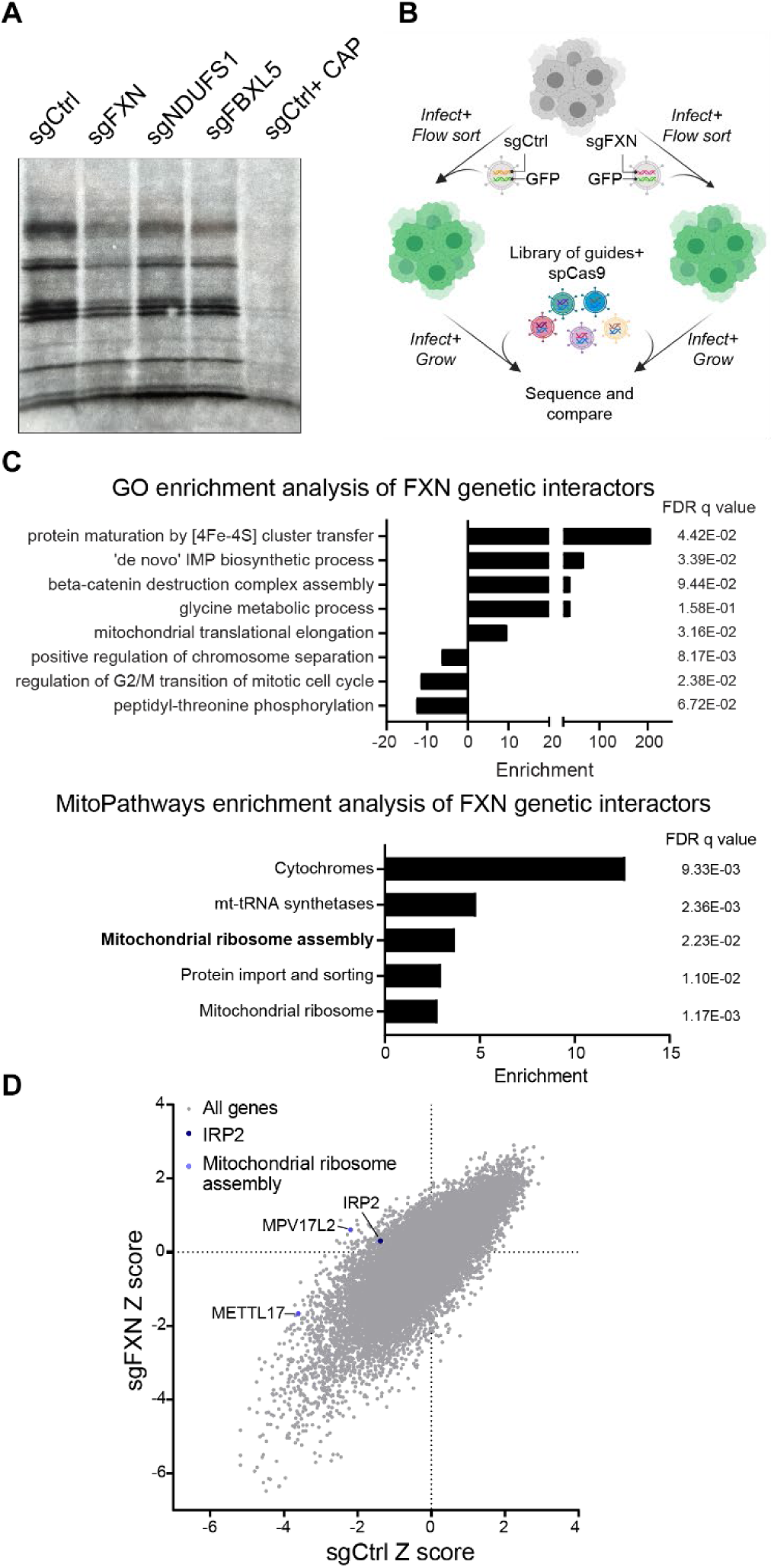
Mitochondrial translation is attenuated in the absence of FXN. A. Mitochondrial translation, as assessed by autoradiography after ^35^S-methionine/cysteine labeling, of cells expressing sgRNAs targeting FXN, NDUFS1 or FBXL5. All cells were treated with 200μg/mL emetine, and control cells in the last lane were also treated with 50μg/μL chloramphenicol. B. Schematic overview of the genome-wide CRISPR genetic interaction screens carried out in K562 cells. Cells were either infected with guides against FXN or a control locus before introduction of the library. Following expansion, cells were sequenced to assess the relative abundance of guides in the FXN null vs. control background. C. GO (top) and MitoCarta 3.0 (bottom) enrichment analysis of genetic interactors identified in for FXN null cells. D. Scatterplot of Z scores showing knockouts growth in sgCtrl vs. sgFXN backgrounds. The positive control (IRP2) and the mitochondrial ribosome assembly genes (METTL17 and MPV17L2) are highlighted.

We wondered which of the many genetic pathways downstream of *FXN* deficiency contribute to the reduced fitness observed in these cells. We performed a genome-wide CRISPR screen (Fig. 2B) on a FXN null background to highlight pathways that become conditionally essential (synthetic sick) vs. redundant or epistatic (buffering) (DeWeirdt et al., 2020). The screen worked from a technical perspective (Fig S2E), and we recovered positive genes / pathways (e.g., *IRP2*; Fig 2D) (Ast et al., 2019) and Fe-S cluster dependent pathways (e.g., *de novo* IMP biosynthesis caused by a loss of PPAT; Fig 2C) in the screen. Intriguingly, enrichment analysis using either GO terms or MitoPathways (from MitoCarta 3.0) highlighted mitochondrial translation as one of the key cellular pathways that was buffered in *FXN* null cells (Fig 2C). Specifically, MitoPathways enrichment highlighted that a key deficit in *FXN* null cells is mitochondrial ribosome assembly, with loss of both the mitoribosome assembly factors MPV17L2 and METTL17 scoring as significant buffering interactions in this screen (Fig 2D and Fig S2F-G). This genetic interaction further highlighted METTL17 as worthy of additional investigation.

### METTL17 is essential to maintain robust mitochondrial translation and is post-transcriptionally depleted in the absence of FXN

METTL17 is a highly conserved mitochondrial matrix protein that has recently been linked to mitochondrial translation (Arroyo et al., 2016). Specifically, METTL17 belongs to the methyltransferase-like family (Wong and Eirin-Lopez, 2021) and has been suggested to take part in the assembly of the mitoribosomal small subunit (Shi *et al*., 2019). However, with respect to the mitoribosome, its methylation target remains controversial (Chen et al., 2020; Itoh et al., 2022; Van Haute et al., 2019), and so the contribution of METTL17 to mitoribosome maturation remains unresolved.

First, we validated our proteomics result that METTL17 is indeed depleted in *FXN* edited cells (Fig 3A, Fig S3A). Moreover, this depletion could not be phenocopied upon loss of *NDUFS1* or *FBXL5*, indicating that it was likely not due to a secondary consequence of OXPHOS deficiency or iron starvation signaling. Importantly, *METTL17* mRNA levels were preserved in *FXN* null cells (Fig 3B), indicating that METTL17 protein is lost post-transcriptionally level in the absence of FXN.

**Fig 3:**
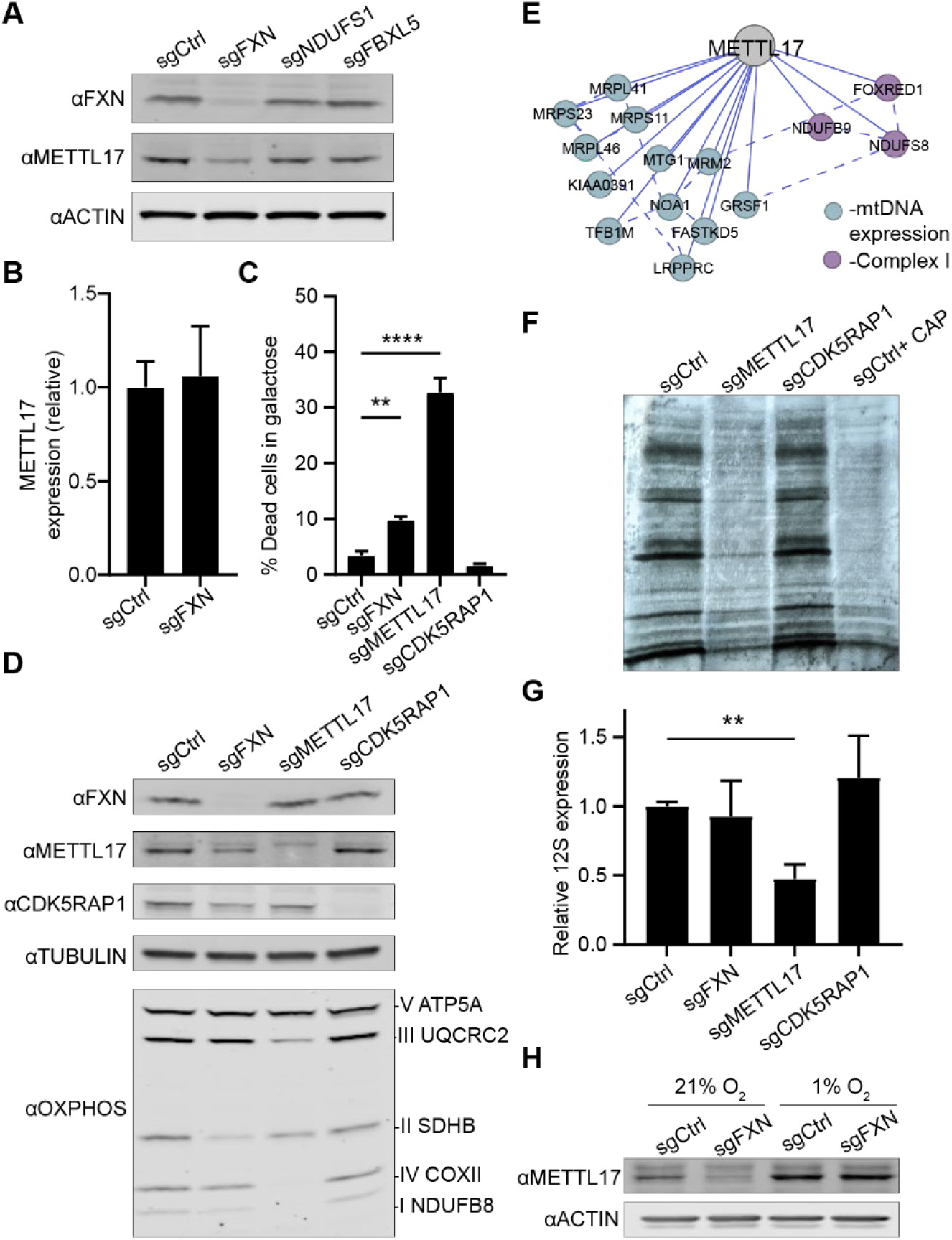
METTL17 is depleted in the absence of FXN and is essential for robust mitochondrial translation. A. Immunoblot for FXN, METTL17 and the loading control actin in K562 cells edited with control, FXN, NDUFS1 and FBXL5 guides. B. qPCR for METTL17 expression levels in sgCtrl and sgFXN cells C. Cells edited for control, FXN, METTL17 and CDK5RAP1 genes were grown for 24h in galactose media, and viability was assessed for each background. D. Immunoblot for FXN, METTL17, CDK5RAP1, select OXPHOS subunits and the loading control tubulin in cells edited with control, FXN, METTL17 and CDK5RAP1 guides. E. Correlation analysis of gene dependencies sourced from DepMap. Presented is the gene network that correlates with METTL17 deletion using FIREWORKS (Amici et al., 2021). Solid and dashed lines represent primary and secondary correlations, respectively. F. Mitochondrial translation, as assessed by autoradiography after ^35^S-methionine/cysteine labeling, of cells expressing sgRNAs targeting METTL17 or CDK5RAP1. All cells were treated with 200μg/mL emetine, and control cells in the last lane were also treated with 50μg/μL chloramphenicol. G. qPCR analysis of 12S levels in cells edited with control, FXN, METTL17 or CDK5RAP1 guides. H. Immunoblot for METTL17 and the loading control actin in cells edited with control or FXN guides. Following editing, cells were grown in 21% or 1% oxygen. All bar plots show mean± SD. **=p < 0.01, ****=p < 0.0001. One-way ANOVA with Bonferroni’s post­test

Next, we sought to determine the extent to which METTL17 depletion attenuates mitochondrial translation and OXPHOS. As with *FXN* depleted cells, *METTL17* null cells undergo extensive cell death when grown on galactose (Fig 3C and Fig S3B), a sugar source that forces cells to rely exclusively on OXPHOS for ATP production (Arroyo *et al*., 2016). No significant galactose-induced death was observed in the absence of CDK5RAP1, a *bona-fide* Fe-S cluster dependent tRNA methylthiotransferase dual localized to the mitochondria and nucleus (Reiter et al., 2012). Furthermore, both *FXN* and *METTL17* null cells displayed combined respiratory chain deficiencies (Fig 3D), in contrast to *CDK5RAP1* depleted cells. METTL17 likely plays a pervasive and crucial role in the maturation of the SSU, consistent with its Cancer Dependency Map profile (Fig 3E), as well as its extensive physical interaction with SSU subunits identified in BioPlex (Fig S3C) (Huttlin et al., 2021). In line with these unbiased methods, we observed near complete ablation of mitochondrial translation and depletion of the 12S (the SSU rRNA) in the absence of METTL17 (Fig 3F-G and Fig S3D). We note that *METTL17* null cells appear to have an even more extreme phenotype than *FXN* edited cells in many of these assays. This may be due to the kinetics of METTL17 depletion in *METTL17* versus *FXN* edited cells, the latter experiencing the METTL17 deficiency as secondary. Finally, we tested whether hypoxia could restore METTL17 levels following loss of *FXN*, as we have previously seen it buffering many FXN-associated phenotypes (Ast *et al*., 2019). Indeed, culturing *FXN* edited cells in 1% O_2_ was sufficient to revert METTL17 levels back to control conditions (Fig 3H), consistent with it being a downstream consequence of FXN deficiency. Collectively, these experiments confirm that METTL17 is indeed intimately linked to SSU functionality in a way that is dependent on intact FXN levels.

### METTL17 contains a putative Fe-S cluster binding motif essential for its function

The striking loss of METTL17 protein levels in the absence of FXN resembles that of *bona-fide* Fe-S cluster containing proteins (Fig 1B) and made us wonder if perhaps METTL17 might also harbor a previously unappreciated Fe-S cluster. Multiple sequence alignment (Fig 4A) of six METTL17 orthologs spanning bacteria, fungi, and humans revealed two striking features: (1) four conserved cysteine residues and (2) a tyrosine residue that forms part of an ‘LYR’ (leucine-tyrosine-arginine) motif in human METTL17. The former is likely a metal binding site, and indeed a recent cryogenic electron microscopy (cryo-EM) structure of the trypanosomal METTL17 homologue revealed coordination of Zn^2+^ through these residues (Saurer et al., 2019). It is known that Zn^2+^ can replace a labile Fe-S cluster in aerobically purified proteins (Imlay, 2006; Nicolet and Fontecilla-Camps, 2014; Wachnowsky and Cowan, 2017). The LYR motif (not to be confused for LYRM protein) is often found in Fe-S cluster apoproteins (Maio et al., 2014) which is proposed to engage the Fe-S cluster handoff machinery. The AlphaFold2.0 predicted structure of human METTL17 (Jumper et al., 2021; Varadi et al., 2022) shows that the conserved cysteines are spatially positioned so as to potentially form an Fe-S binding pocket while the LYR motif is surface accessible, further strengthening the likelihood that these motifs are functional. Moreover, METTL17 remains stable following disruption of the cytosolic Fe-S biosynthesis machinery (Fig S4A), further hinting at a specific connection between loss of METTL17 and mitochondrial Fe-S cluster assembly. Collectively, these lines of evidence led us to hypothesize that METTL17 is a mitochondrial Fe-S cluster binding apoprotein, and that perhaps this was previously missed given the labile nature of Fe-S clusters.

**Fig 4:**
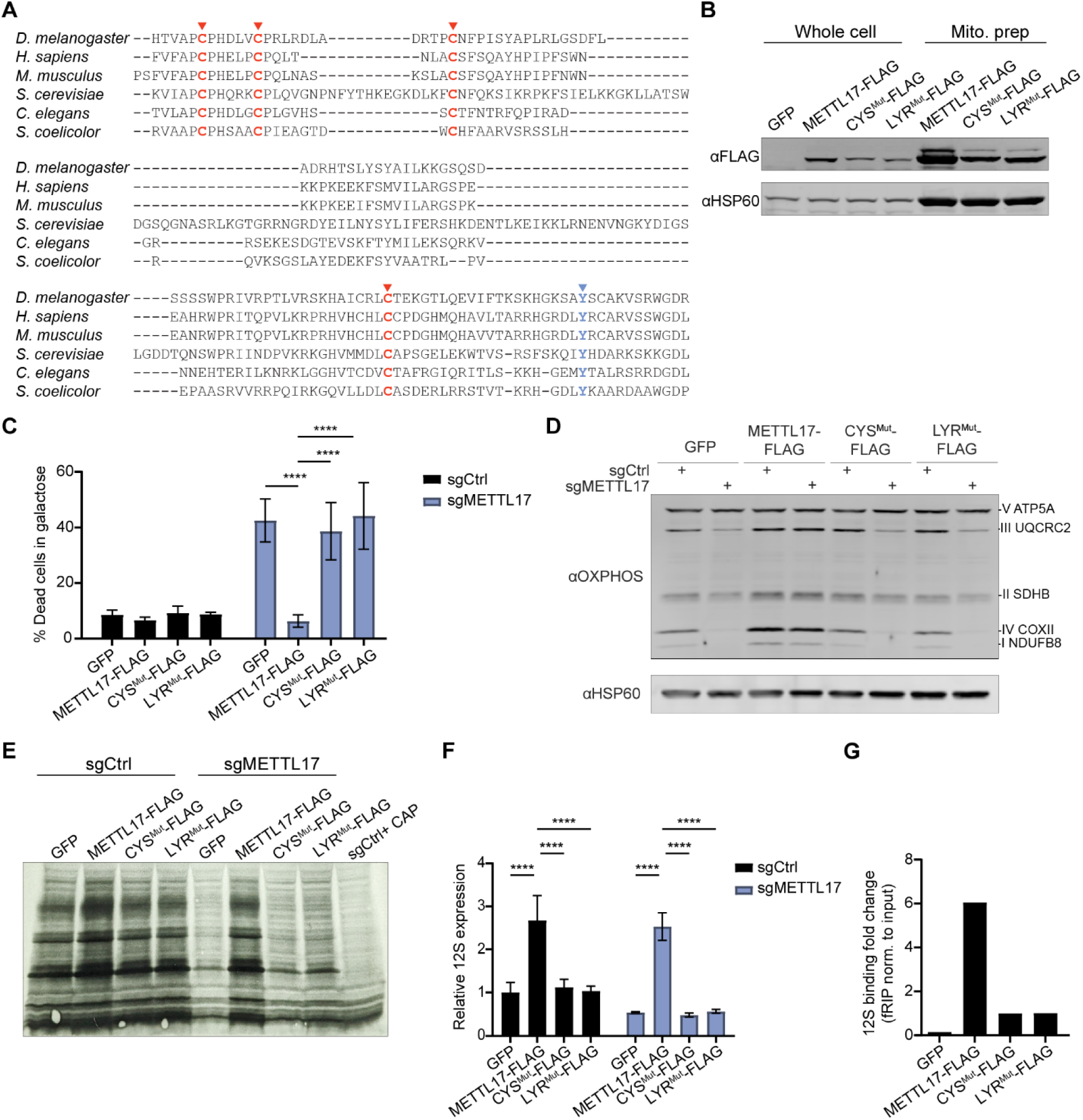
METTL17 has two conserved motifs linked to Fe-S binding, which are crucial for its functionality. A. Multiple sequence alignment for METTL17 homologues, highlighting two motifs associated with Fe-S cluster binding; 4 cysteine metal binding pocket (red) and a LYR handoff motif (blue). B. Immunoblot from whole cell and mitoprep extracts of cells expressing GFP, METTL17-FLAG, CYS^Mut^-FLAG or LYR^Mut^-FLAG constructs. C. Control or METTL17 edited cells expressing GFP, METTL17-FLAG, CYS^Mut^-FLAG or LYR^Mut^-FLAG constructs were grown for 24h in galactose, following which their viability was assessed. D. Immunoblots examining OXPHOS subunits or the loading control HSP60 in Control or METTL17 edited cells expressing GFP, METTL17-FLAG, CYS^Mut^-FLAG or LYR^Mut^-FLAG constructs. E. Mitochondrial translation, as assessed by autoradiography after ^35^S-methionine/cysteine labeling, in Control or METTL17 edited cells expressing GFP, METTL17-FLAG, CYS^Mut^-FLAG or LYR^Mut^-FLAG constructs. All cells were treated with 200μg/mL emetine, and control cells in the last lane were also treated with 50μg/μL chloramphenicol. F. qPCR analysis of 12S levels in Control or METTL17 edited cells expressing GFP, METTL17-FLAG, CYS^Mut^-FLAG or LYR^Mut^-FLAG constructs. G. Formaldehyde-linked RNA immunoprecipitation of the 12S to GFP, METTL17-FLAG, CYS^Mut^-FLAG or LYR^Mut^-FLAG proteins. Results were normalized to input construct and 12S levels. All bar plots show mean ± SD. ****=p < 0.0001. One-way ANOVA with Bonferroni’s post-test.

We asked whether these candidate Fe-S binding motifs were indeed important for the activity of METTL17. To this end, we generated three constructs for METTL17, a WT FLAG-tagged version as well as two mutants: CYS^Mut^-FLAG in which the conserved cysteines were changed to serines, and LYR^Mut^-FLAG in which the LYR residues were mutated to alanines. All three of these variant proteins were enriched in mitochondrial fractions (Fig 4B), demonstrating their proper subcellular targeting, although the two mutants were less abundant than the WT (as is typical of Fe-S cluster apoproteins lacking their cofactor). We next tested if these mutants could support mitochondrial translation in the absence of endogenous METTL17. METTL17-FLAG expression could rescue for loss of the endogenous gene as shown by reduced death in galactose and restored OXPHOS subunit levels, mitochondrial de-novo translation and 12S levels (Fig 4C-F and Fig SB-C). However, the CYS^Mut^-FLAG and LYR^Mut^-FLAG mutants failed to rescue any of the defects assayed. This would indicate that both motifs are crucial for METTL17’s role in supporting SSU activity. Of note, METTL17-FLAG expression served to boost mitochondrial translation and the steady state levels of multiple OXPHOS subunits (most notably in complexes I, II and IV) even above the GFP control, suggesting that it is a rate limiting factor in mitochondrial protein synthesis.

We next wondered whether the cysteine binding pocket and LYR motif were also required for METTL17 engagement with the SSU. We performed a formaldehyde RNA immunoprecipitation assay (D et al., 2016) to test for the interaction between the various constructs and the 12S (Fig 4G). In this assay, cells are lightly crosslinked, lysed, the protein of interest is immunoprecipitated and bound RNA is quantified by qPCR. We could observe that while METTL17-FLAG showed robust binding to the 12*S* as compared to a GFP control, the interaction between CYS^Mut^-FLAG or LYR^Mut^-FLAG and the 12*S* was diminished (even taking into account the lower basal expression levels for the constructs compared with 12*S*). In contrast, there was only marginal enrichment for 16*S* binding for these constructs when compared to the GFP control, and no notable difference between the WT and mutant forms of METTL17 (Fig S4D). These findings suggest that apart from structural stability, the putative Fe-S cluster in METTL17 is also important for contributing (directly or indirectly) to SSU and 12*S* binding.

### Biophysical studies confirm METTL17 harbors an Fe-S cluster

We then set out to study and analyze the purified human METTL17 under conditions that would be optimal to preserve any Fe-S cluster present. Affinity chromatography of METTL17 gave a clean preparation of the protein that resulted in both a single band on an SDS-PAGE gel and a monodisperse analytical gel filtration peak corresponding to a molecular weight near 50 kDa (Fig 5A). Purified METTL17 precipitates upon exposure to air as is common for air-sensitive Fe-S proteins (Wachnowsky and Cowan, 2017). Inductively coupled plasma mass spectrometry (ICP-MS) analysis revealed the presence of 3.7 Fe ions per polypeptide (Fig 5B). In a variant in which the four cysteines predicted to coordinate the cluster (C333, C339, C347, and C404) were mutated to serines (‘CYS^Mut^’), almost all the iron was lost (0.2 Fe ions per polypeptide). The UV-Vis spectrum of METTL17 contains a broad charge-transfer band around 400 nm, suggesting the presence of either an [Fe_4_S_4_]^2+^ cluster or an [Fe_3_S_4_]^+^ cluster (Fig. 5C) (Agar et al., 2000; Bruschi et al., 1976; Emptage et al., 1983; Khoroshilova et al., 1997); this band is absent in the spectrum of the CYS^Mut^ sample. No electron paramagnetic resonance (EPR) signal was observed for the METTL17 sample between 5 and 50 K (data not shown), and addition of the reducing agent dithionite or the oxidizing agent indigodisulfonic acid also did not produce an EPR signal. The lack of an EPR signal in the as-isolated and oxidized samples rules out the presence of an [Fe_3_S_4_]^+^ cluster, which has a characteristic EPR signal (Hagen, 1992). Taken together, the Fe quantitation, UV-Vis spectroscopy, and EPR spectroscopy results are most consistent with the presence of an [Fe_4_S_4_]^2+^ cluster coordinated by C333, C339, C347, and C404 on human METTL17.

**Fig 5:**
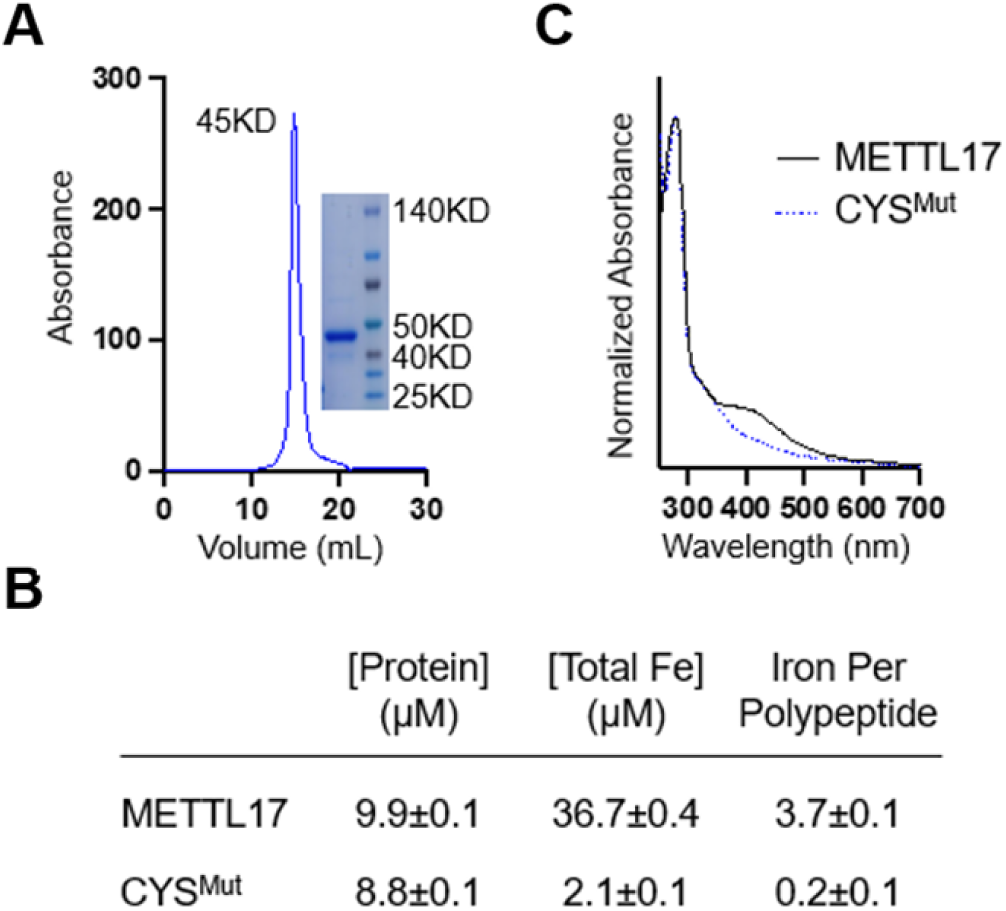
Human METTL17 expressed and purified from *E. coli* contains an Fe-S cluster. A. Gel filtration chromatography and SDS-PAGE analysis demonstrate that the purified METTL17 construct runs as a monomer near its predicted molecular weight of 50 kD. B. Iron content of purified METTL17and CYS^Mut^ as determined by bicinchoninic acid assay and inductively coupled plasma mass spectrometry. The CYS^Mut^ is a METTL17 variant in which the four cysteines predicted to coordinate the cluster are mutated to serine (C333S, C339S, C347S, and C404S). C. The UV-Vis absorption spectra of METTL17 exhibits a broad band around 420 nm, consistent with the presence of an [Fe_4_S_4_]^2+^ or an [Fe_3_S_4_]^+^ cluster; the latter is ruled out by EPR spectroscopy as described in the text. This band is lost in the spectrum of CYS^Mut^. For clarity, spectra were normalized to the intensity at 280 nm.

### The [Fe_4_S_4_]^2+^ cluster on METTL17 stabilizes its binding to the mitoribosomal small subunit prior to translation initiation

Next, we sought to directly visualize the Fe-S cluster in METTL17 using cryo-EM, with a focus on capturing it on the mitoribosome. We made initial attempts to study the human SSU-METTL17 complex, but METTL17 escaped detection as its binding is likely not stable enough. As the putative Fe-S cluster binding motif is conserved (Fig 4A), we used mitoribosomes from the yeast *S. cerevisiae*, which provide a suitable model for structural studies due to stabilising rRNA expansion segments and protein extensions (Amunts et al., 2014) that would be amenable for studying assembly intermediates. Although previous fungi mitoribosome structures have been reported (Desai et al., 2017; Itoh et al., 2020), they did not investigate SSU assembly.

Reconstructions from the first cryo-EM dataset followed by 3D classification and refinement revealed two distinct classes, the empty and METTL17-bound states of the SSU (Fig S5). In the second dataset, we added a recombinant initiation factor mtIF3, and obtained three states: free SSU, METTL17-bound, mtIF3-bound. We then grouped particles from the two datasets and resolved the structures of the SSU, SSU-METTL17, and SSU-mtIF3 at an overall resolution of 2.3 Å, 2.6 Å, and 2.8 Å, respectively (Table S1). Compared to the previous report of *S. cerevisiae* SSU at 3.5 Å resolution (Desai *et al*., 2017), the improved reconstruction allowed us to build a more accurate model. Specifically, we modelled extensions of the mitoribosomal proteins bS1m, uS2m, uS3m and uS7m as well as corrected the assignment of a guanosine diphosphate (GDP) to adenosine triphosphate (ATP) in the nucleotide pocket of the protein mS29 (S6 A-D).

In the SSU-METTL17 complex, the 72-kDa factor is found in the cleft between the head and the body and its presence prevents a monosome formation (Fig 6A). In contrast to previously characterised SSU assembly factors (Itoh *et al*., 2022), METTL17 binds exclusively to the rRNA of the head, and the binding involves both, the N-terminal domain (NTD) and the C-terminal domain (CTD). The NTD is a Rossmann-fold methyltransferase domain, characterized by a seven-stranded beta-sheet core sandwiched by six alpha-helices, while the CTD is a unique fold that consists of a four-stranded beta-sheet with an alpha-helix. The comparison with the non-bound state shows that the association of METTL17 withdraws the rRNA helix h31 causing rearrangement of h32 and h34, and the entire head is rotated by approximately 3 Å up to open the tRNA binding cleft. Thus, h31 is more exposed, and the rRNA nucleotide A1100 in h34 is flipped out (Fig. 6B). In addition, the mitochondrial C-terminal extension fills the mRNA channel, blocking its premature binding.

**Fig 6:**
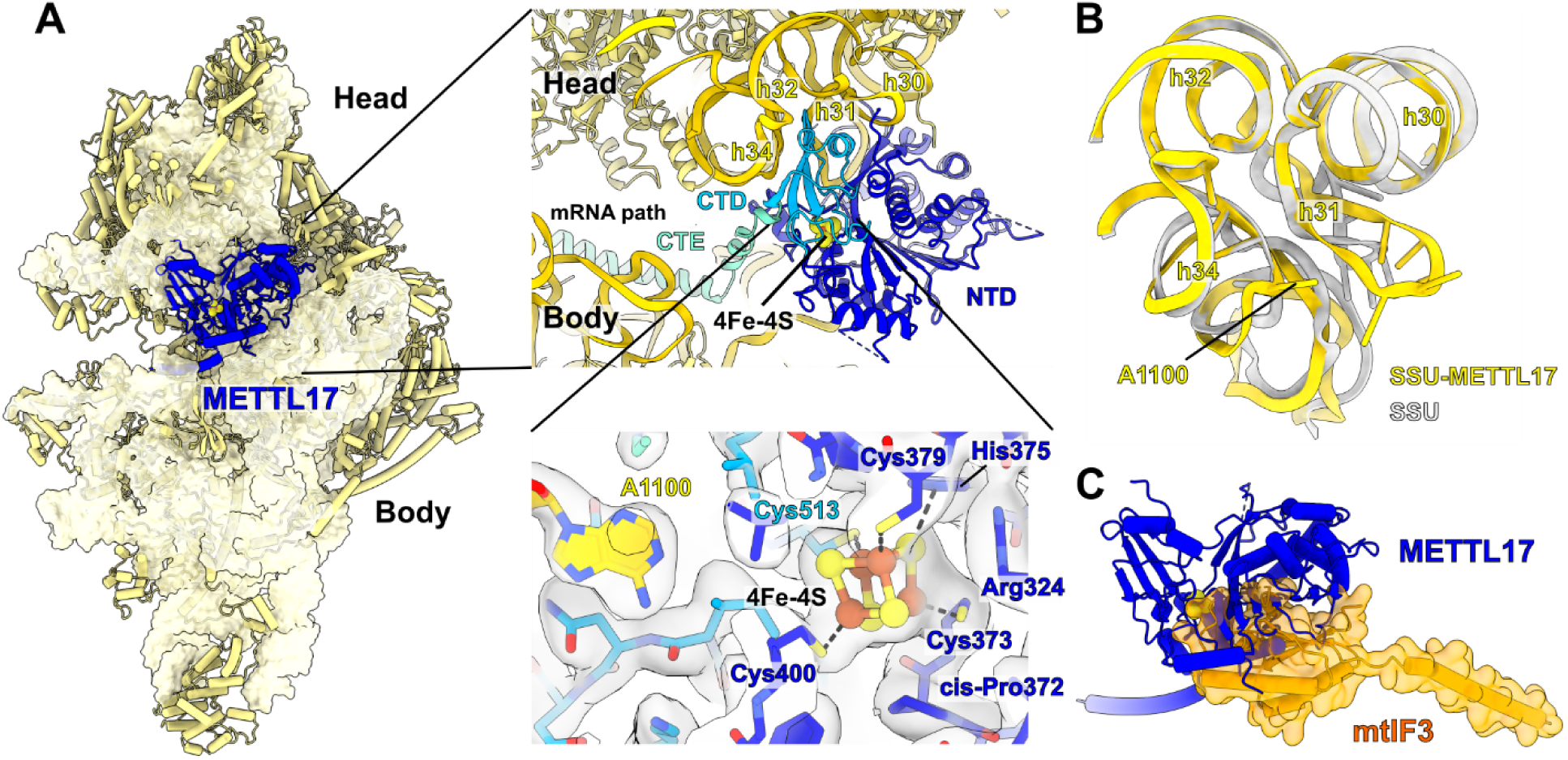
Cryo-EM structure of the yeast SSU-METTL17 complex and involved elements. A. Overall view of METTL17 on the SSU, and close-up views. Top close up shows the position of METTL17 (C-terminal domain, CTD light blue; N-terminal domain, NTD blue) between the rRNA (yellow) of the head and body, while the C-terminal extension (CTE) occupies the mRNA path. Bottom close up shows the coordination of 4Fe-4S cluster by four cysteines, including Cys513 from the CTD, and related structural elements with their cryo-EM densities: flipped base A1100, a cis-proline, arginine that is within salt bridge distance, and a conserved histidine that can be involved in a transfer and ligation to the Fe-S unit. B. Conformational changes within the rRNA region h30-34 that is involved in METTL17 binding. Superimposed models of the SSU-METTL17 (yellow) with unbound state (grey). Sticks represent the rRNA residues responsible for the METTL17 interaction. C. Superposition of SSU-METTL17 with SSU-mtIF3 showing clashes of METTL17 (blue) with mtIF3 (orange surface representation).

The SAM motif Gly-X-Gly-X-Gly of *S. cerevisiae* METTL17 is disrupted with an Ala at the last Gly position and a Tyr at the first X position, suggesting that the methylation is not a conserved function of this protein in the process of the mitoribosome assembly. In fact, the 15S SSU rRNA in yeast is not methylated (Klootwijk et al., 1975) and no electron density was observed for a SAM cofactor. Of note, a distance of 25 Å separates the metal binding site from the SAM binding motif in the cryo-EM structure of the trypanosomal METTL17 homologue (Saurer *et al*., 2019), arguing against METTL17 functioning as a radical SAM enzyme in this organism as well. We identified an ordered density, adjacent to sulfhydryl groups of four cysteine residues Cys373, Cys379, Cys400, Cys513 (yeast numbering, equivalent to human Cys333, Cys339, Cys347, Cys404) (Fig 6A). The density corresponds to eight atoms organized in a cube, which is consistent with [Fe_4_S_4_]^2+^ cluster, where the four iron atoms are bound by cysteines S atoms and bridge sulfur atoms. The unique feature of the embedded [Fe_4_S_4_]^2+^ cluster in our structure is that it brings together residues that are 140 amino acids apart, thus stabilising the N- and C-terminal domains of METTL17 when it’s bound to the SSU. The resulting orientation of Cys400 and Cys513 forms a pocket, where rRNA base A1100 adapts, which contributes to a prominent binding of all the components together on the mitoribosome. Moreover, we identified a cis-proline Pro372 of METTL17 that orients the Cys373 side chain towards the [Fe_4_S_4_]^2+^ cluster and further supports the binding. Furthermore, we identified His375 and Arg324 within cluster interaction distance, suggesting a potential involvement in transfer and ligation of the [Fe_4_S_4_]^2+^ cluster. The AlphaFold2.0 predicted structure of the human METTL17 (Jumper *et al*., 2021; Varadi *et al*., 2022) is overall consistent with our model (Fig S6E), and the superposition with the high resolution structure of the human SSU (Itoh *et al*., 2022)suggests conserved interacting interfaces (Fig S6F).

To further clarify a putative role of METTL17, we formed a preinitiation complex by purifying the SSU-METTL17 and incubating it with a recombinant mtIF3. A cryo-EM density map was then calculated, and we identified a new class containing a subset of 53,922 particles that was processed with a signal subtraction, followed by 3D classification using the mask for the mtIF3-binding site (Fig S5). The map features the complex SSU-mtIF3, where the initiation factor occupies a position on the SSU that is similar to the human counterpart (Fig S6G) (Khawaja et al., 2020) and mutually exclusive with METTL17, as a superposition of the two states shows clashes between the factors (Fig 6C). Thus, the binding of the regulatory factor METTL17 is stabilised by the [Fe_4_S_4_]^2+^ cluster and precedes the initiation of translation.

### Forced expression of METTL17 rescues bioenergetic, but not growth, defects of FXN null cells

We sought to determine which of the downstream phenotypic consequences of FXN deficiency could be prevented by forced expression of METTL17. We tested the effects of overexpressing METTL17, or its mutants, in *FXN* null cells. Remarkably, we observed that the galactose-induced death of *FXN* null cells could be reversed by the forced overexpression of METTL17-FLAG, but not CYS^Mut^-FLAG or LYR^Mut^-FLAG (Fig 7A). In line with these findings, we showed that the loss of various OXPHOS protein subunits and the lowered oxygen consumption of FXN null cells were restored to WT levels by overexpression of METTL17-FLAG (Fig 7B-C and Fig S7A-C). Thus, it seems that the bioenergetic defect of *FXN* depleted cells could largely be restored by re-expressing one mitochondrial Fe-S cluster containing protein: METTL17.

**Fig 7:**
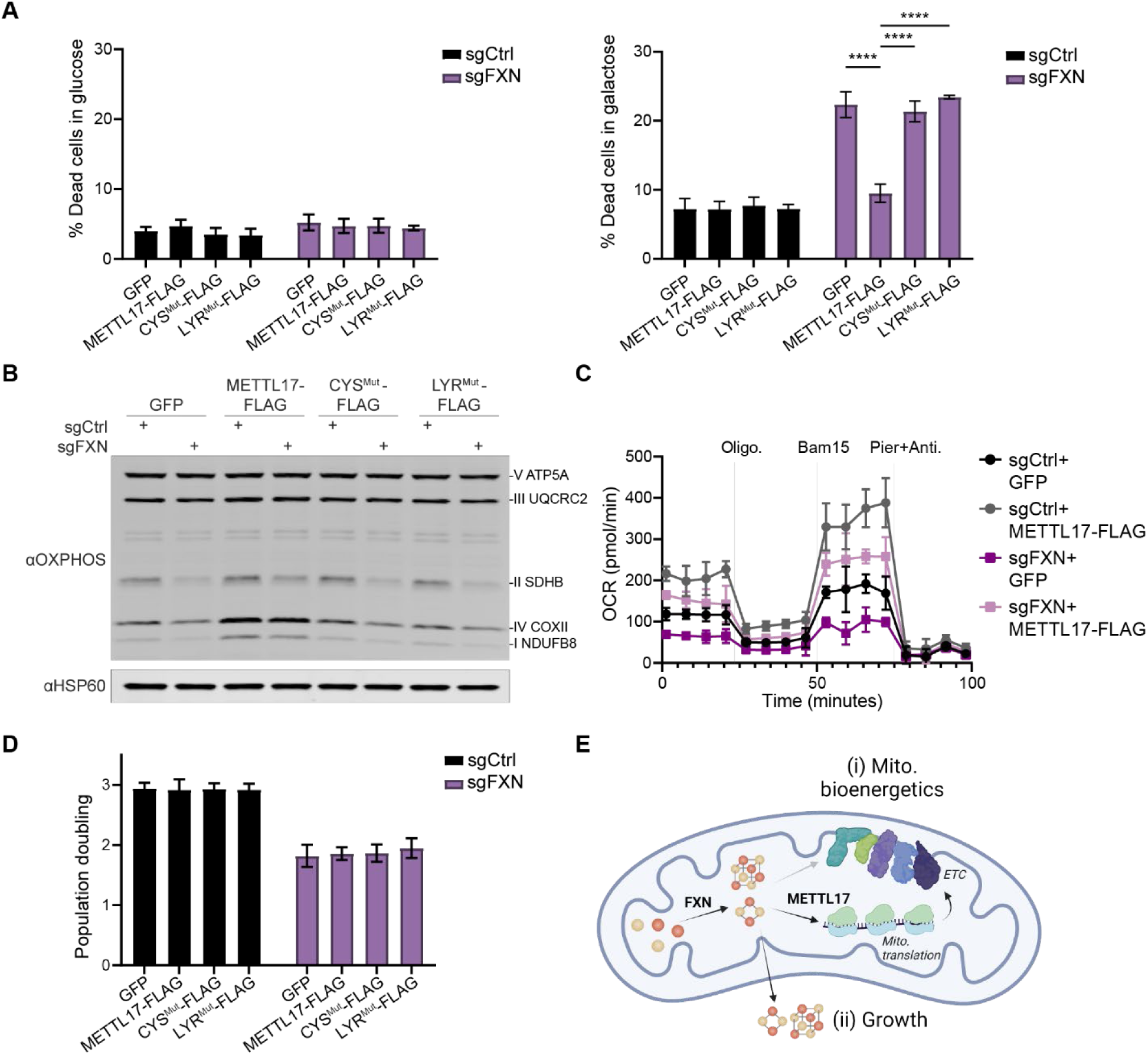
Overexpression of METTL17 restores the mitochondrial bioenergetics, but not growth, of FXN null human cells. A. Control or FXN edited cells expressing GFP, METTL17-FLAG, CYS^Mut^-FLAG or LYR^Mut^-FLAG constructs were grown for 24h in glucose (left) or galactose (right), following which their viability was assessed. B. Immunoblots examining OXPHOS subunits or the loading control HSP60 in Control or FXN edited cells expressing GFP, METTL17-FLAG, CYS^Mut^-FLAG or LYR^Mut^-FLAG constructs. C. Oxygen consumption rate (OCR) of Control or FXN edited cells expressing GFP or METTL17-FLAG. Cells were sequentially treated with oligomycin, Bam15 and piericidin+antimycin. D. Population doubling over 72h of Control or FXN edited cells expressing GFP, METTL17-FLAG, CYS^Mut^-FLAG or LYR^Mut^-FLAG constructs. E. FXN activates Fe-S cluster formation, which can be utilized to support (i) mitochondrial bioenergetics via formation of the electron transport chain (ETC) or (ii) cell growth and division. METTL17 is a key Fe-S cluster bearing modulator of mitochondrial bioenergetics, and its absence in FXN null cells accounts for much of the mito. bioenergic defects observed in these cells. All bar plots show mean± SD. **=p < 0.01, ****=p < 0.0001. One-way ANOVA with Bonferroni’s post-test.

Next, we wondered if other cellular defects outside of mitochondria and observed in FXN null cells could be rescued by METTL17 overexpression, as human Fe-S cluster biosynthesis is required for other essential processes such as nuclear DNA maintenance, repair and nucleotide synthesis. When we tested the growth of *FXN* null cells upon METTL17 overexpression, however, we saw no benefit (Fig 7D). Moreover, the cytosolic Fe-S apoprotein POLD1 remained depleted in *FXN* null cells irrespective of METTL17 overexpression (Fig S7D).

Together, these data demonstrate for the first time that the growth and bioenergetic defects observed in *FXN* null cells are separable, and that they can be uncoupled from each other through the expression of the Fe-S cluster containing protein METTL17. Fe-S clusters that are made in the mitochondria can in principle be directed to two distinct routes: one for local use within mitochondria supporting bioenergetic demand vs. one for export to the cytosol to support growth (Fig 7E). METTL17 expression can evidently privilege Fe-S clusters for use within mitochondria.

## Discussion

Here we have made the discovery that METTL17, a previously described putative SSU maturation factor, harbors a hitherto unrecognized [Fe_4_S_4_] cluster that is critical for its function in protein synthesis. We report for the first time that METTL17 deficiency lies downstream of *FXN* loss and appears to be a key effector of impaired mitochondrial translation. Re-expressing METTL17 is sufficient to rescue the bioenergetic, but not growth, phenotypes associated with *FXN* deficiency. These findings demonstrate that the Fe-S cluster supply within mitochondria can be uncoupled from that of the cytosol, and that factors such as METTL17 can privilege the availability of Fe-S clusters for specific routes.

Our proteome-wide analysis indicates that nearly all the Fe-S cluster containing proteins we could detect are de-stabilized when the Fe-S cluster supply is reduced. While it has long been known that one function of Fe-S clusters (in addition to their many roles in electron transfer, oxyanion binding, etc.) is protein stability (Lill and Freibert, 2020), our findings show that in fact this is a nearly universal role of the cluster. Previous proteomic profiling of FA patient cells has described deficits in some, but not all, Fe-S cluster containing proteins (Telot et al., 2018). However, it is likely that a near total depletion of *FXN*, such as is generated by CRISPR editing is not observed in patients.

Our structural studies have revealed that the [Fe_4_S_4_]^2+^ cluster stabilizes METTL17 and assists in its coupling to the rRNA of the mitoribosomal small subunit during the assembly process. Throughout evolution, the presence of an [Fe_4_S_4_]^2+^ cluster on METTL17 is more conserved than that of SAM binding. When examining the METTL17 ortholog *D. discoideum*, a member of the Amoebozoa outgroup of metazoa and fungi, we find that both the SAM motif and the four cysteine residues coordinating Fe-S cluster binding are present indicating that this is likely the more ancient form of the protein. The obtained and predicted structure of METTL17 argues against a radical SAM role for the Fe-S, given its distance to the SAM binding motif and the fact that the iron sulfur cluster is coordinated by four cysteines rather than three. The obtained structure of the SSU-METTL17 complex allows us to put METTL17 in the context of the dynamic SSU assembly, extending its part in the view of the mitoribosome formation. Comparison with structural studies of earlier assembly intermediate states (Itoh *et al*., 2022) suggests that METTL17 can be accommodated on the SSU in the presence of RBFA, and we also show that METTL17 is replaced by the initiation factor mtIF3. Thereby, METTL17 facilitates a productive assembly pathway through acting on relatively late stages, which reflects the structural importance of the [Fe_4_S_4_]^2+^ bound METTL17 for the biological function of the mitoribosome biogenesis.

Our data indicates that METTL17 operates as an Fe-S cluster “checkpoint” for protein synthesis. The mitochondrial respiratory chain is rich with Fe-S clusters, and our work shows that mitochondrial translation is dependent on METTL17 being present with an intact Fe-S cluster for proper ribosome assembly. We have shown that METTL17 is very labile in the absence of Fe-S levels, and moreover, that METTL17 is limiting for mitochondrial translation. Hence, METTL17 can ensure that mitochondrial translation proceeds only when Fe-S cluster levels are replete. As solvent exposed Fe-S clusters are extremely labile to reactive nitrogen and oxygen species, METTL17 could serve as a “kill switch” for turning off the production of mitochondrial machinery in unfavorable environments, enabling sub-organelle regulation of mitochondrial protein synthesis to match local redox conditions (Allen, 2015).

Our findings indicate that at least in cultured cells, *FXN* deficiency results in a previously under-appreciated deficit in mitochondrial protein synthesis, which appears to be largely downstream of METTL17. Although OXPHOS deficiency, specifically loss of CI and CII, have long been appreciated to be features of FA samples (Puccio *et al*., 2001; Rotig *et al*., 1997), to the best of our knowledge, no prior report has documented defects in mitochondrial protein synthesis. Future efforts should be aimed at determining whether patient derived specimens show any evidence of impaired mitochondrial translation. Recent work has identified Fe-S cluster proteins involved in mitochondrial translation embedded in tRNA maturation (Reiter *et al*., 2012; Wei et al., 2015) as well as in the SSU (Itoh *et al*., 2022). However, overexpression of METTL17 seems to be sufficient to restore the bioenergetic defects of FXN null cells, suggesting it may be limiting for mitochondrial protein synthesis. This property of METTL17 is unusual and suggests it could represent an interesting therapeutic target, not only for FA. In principle, other inherited OXPHOS disorders-independent of genetic etiology-may also benefit from boosting mitochondrial protein synthesis via METTL17 overexpression. Further experiments will be required to test this hypothesis.

A curious feature of METTL17 is that its expression uncouples the mitochondrial versus cytosolic routing of Fe-S clusters. In animal cells, *de novo* Fe-S cluster biosynthesis begins inside of mitochondria, and this supply is then conveyed within the organelle and also exported to the cytosol (Lill and Freibert, 2020; Maio and Rouault, 2020; Srour *et al*., 2020). Simply forcing the expression of METTL17 seems to privilege mitochondrial-generated Fe-S clusters for intra-mitochondrial use. It is interesting to speculate whether additional, as yet unidentified factors prioritize cytosolic routing of Fe-S clusters. The ability to uncouple these routes by expressing METTL17 could be useful for dissecting the pathogenesis of FA, since at present, we do not know which is dominant in the etiopathogenesis: bioenergetic defect or growth defect. If the bioenergetic defects predominate *in vivo*, boosting METTL17 could hold therapeutic potential.

## ACKNOWLEDGMENTS

We wish to thank T.L. To, Timothy Durham, and John G. Doench for technical assistance, and all members of the Mootha lab for fruitful discussions and feedback. This work was supported by the Friedreich’s Ataxia Research Alliance (FARA), the European Research Council (ERC-2018-StG-805230), and the Knut and Alice Wallenberg Foundation (2018.0080). D.L.M.S. acknowledges funding from the National Institutes of Health under award number GM141203. Y.I. was supported by H2020-MSCA-IF-2017 (799399-Itohribo). Support for the ICP-MS instrument was provided by a core center grant P30-ES002109 from the National Institute of Environmental Health Sciences, NIH. The SciLifeLab cryo-EM facility is funded by the Knut and Alice Wallenberg, Family Erling Persson, and Kempe foundations. V.K.M. is an Investigator of the Howard Hughes Medical Institute.

## Author Contributions

TA, YI, SS, JGM, PRJ, IC, PK DLMS, AA, VKM designed research; TA, YI, SS, JGM, GN, PRJ, PK, IC performed research and analyzed data; DLMS, PK, AA and VKM supervised the study; TA and VKM wrote the manuscript with input from all authors.

## Conflicts of Interest

VKM is on the scientific advisory boards of Janssen Pharmaceuticals and 5AM Ventures. TA and VKM are listed as inventors on a patent application filed by The Broad Institute on the use of METTL17 as a therapy.

## Methods

### Cell Lines

K562 (female), HEK293T (female) and A549 (male) cells were obtained from the ATCC and maintained in DMEM (GIBCO) with 25 mM glucose, 10% fetal bovine serum (FBS, Invitrogen), 4mM Glutamine, 1 mM sodium pyruvate, 50 µg/mL uridine, and 100 U/mL penicillin/ streptomycin under 5% CO_2_ at 37°C. When necessary, K562 cells were selected with 2 µg/ml puromycin (GIBCO) or 500ug/ml Geneticin (GIBCO), HEK293T and A549 cells were selected with 1 µg/ml puromycin (GIBCO). Cell lines were authenticated by STR profiling (ATCC). Cells were tested to ensure absence of mycoplasma by PCR-based assay once every 3 months. For experiments involving 1% oxygen, cells were placed in 37°C incubators, attached to a nitrogen supply which pulsed N2 and maintained in 1% O_2_ and 5% CO_2._

### Plasmids

Individual sgRNAs were cloned into pLentiCRISPRv2 (Addgene 52961) (Sanjana et al., 2014). For genetic interaction assays, cells were infected with pRDA_186 plasmid (Addgene 133458), a gift from John Doench (Broad Institute), bearing guides against a control locus or FXN. C-terminally FLAG tagged, codon optimized cDNAs were cloned in pLYS6, bearing a Neomycin selection cassette, using the NheI and EcoRI sites. All plasmids were verified by sequencing. pMD2.G (Addgene 12259) and psPAX2 (Addgene 12260) were used for lentiviral packaging.

### Lentivirus production

2.5×10^6^ HEK293T cells were seeded in 5ml in a T25cm^2^ flask (one flask per lentivirus). The following day the cells were transfected with 1ml of transfection mixture per well. The transfection mixture contained 25 µl Lipofectamine 2000 (Thermo Fisher Scientific), 3.75µg psPAX2, 2.5µg pMD2.G, 5µg of the lentiviral vector of interest and Opti-MEM medium (GIBCO) up to 1ml. The mixture was incubated at room temperature for 20 min before adding it to cells. 6h following transfection, the media was replaced with fresh DMEM. Two days after transfection, media was collected, filtered through a 0.45um filter and stored at –80C.

### Proteomics

sgCtrl and sgFXN K562 cells were grown in duplicate flasks for 10 days. Quantitative proteomics was performed at the Thermo Fisher Scientific Center for Multiplexed Proteomics (Harvard).

#### Sample Preparation for Mass Spectrometry

Samples were prepared essentially as previously described (Li et al., 2021; Navarrete-Perea et al., 2018). Following lysis, protein precipitation, reduction/alkylation and digestion, peptides were quantified by micro-BCA assay and 100µg of peptide per sample were labeled with TMT reagents (Thermo-Fisher) for 2hrs at room temperature. Labeling reactions were quenched with 0.5% hydroxylamine and acidified with TFA. Acidified peptides were combined and desalted by Sep-Pak (Waters).

#### Basic pH reversed-phase separation (BPRP)

TMT labeled peptides were solubilized in 5% ACN/10 mM ammonium bicarbonate, pH 8.0 and separated by an Agilent 300 Extend C18 column (3.5µm particles, 4.6 mm ID and 250 mm in length). An Agilent 1260 binary pump coupled with a photodiode array (PDA) detector (Thermo Scientific) was used to separate the peptides. A 45 minute linear gradient from 10% to 40% acetonitrile in 10 mM ammonium bicarbonate pH 8.0 (flow rate of 0.6 mL/min) separated the peptide mixtures into a total of 96 fractions (36 seconds). A total of 96 Fractions were consolidated into 24 samples in a checkerboard fashion, acidified with 20 µL of 10% formic acid and vacuum dried to completion. Each sample was desalted via Stage Tips and re-dissolved in 5% FA/ 5% ACN for LC-MS3 analysis.

#### Liquid chromatography separation and tandem mass spectrometry (LC-MS3)

Proteome data were collected on an Orbitrap Eclipse mass spectrometer (ThermoFisher Scientific) coupled to a Proxeon EASY-nLC 1200 LC pump (ThermoFisher Scientific). Fractionated peptides were separated using a 180 min gradient at 500 nL/min on a 35 cm column (i.d. 100 μm, Accucore, 2.6 μm, 150 Å) packed in-house. MS1 data were collected in the Orbitrap (120,000 resolution; maximum injection time 50 ms; AGC 4 × 105). Top 10 precursors with charge states between 2 and 5 were required for MS2 analysis, and a 90 s dynamic exclusion window was used. MS2 scans were performed in the ion trap with CID fragmentation (isolation window 0.5 Da; Rapid; NCE 35%; maximum injection time 35 ms; AGC 1.5 × 104). An on-line real-time search algorithm (Orbiter) was used to trigger MS3 scans for quantification (Schweppe et al., 2020). MS3 scans were collected in the Orbitrap using a resolution of 50,000, NCE of 55%, maximum injection time of 150 ms, and AGC of 1.5 × 105. The close out was set at two peptides per protein per fraction(Schweppe *et al*., 2020).

#### Data analysis

Raw files were converted to mzXML, and monoisotopic peaks were re-assigned using Monocle (Rad et al., 2021). Searches were performed using SEQUEST (Eng et al., 1994) against a human database downloaded from Uniprot in 2014. We used a 50 ppm precursor ion tolerance and 0.9 Da product ion tolerance for MS2 scans collected in the ion. TMT on lysine residues and peptide N-termini (+229.1629 Da) and carbamidomethylation of cysteine residues (+57.0215 Da) were set as static modifications, while oxidation of methionine residues (+15.9949 Da) was set as a variable modification.

Each run was filtered separately to 1% False Discovery Rate (FDR) on the peptide-spectrum match (PSM) level. Then proteins were filtered to the target 1% FDR level across the entire combined data set. For reporter ion quantification, a 0.003 Da window around the theoretical m/z of each reporter ion was scanned, and the most intense m/z was used. Reporter ion intensities were adjusted to correct for isotopic impurities of the different TMT reagents according to manufacturer specifications. Proteins were filtered to include only those with a summed signal-to-noise (SN) ≥ 100 across all TMT channels. For each protein, the filtered peptide TMT SN values were summed to generate protein quantification values. To control for different total protein loading within a TMT experiment, the summed protein quantities of each channel were adjusted to be equal within the experiment.

### Mito translation assay

2.5×10^6^ cells expressing the corresponding sgRNAs were washed in PBS and incubated for 30 min in 1 mL of labeling medium (10% dFBS, 1 mM sodium pyruvate, and 50 μg/mL uridine in DMEM without methionine/cysteine; Life Technologies). Emetine (Sigma) was added to a final concentration of 200 μg/mL, and cells were incubated for 5 min before addition of 200μCi 35S-labeled methionine/cysteine mixture (PerkinElmer) and incubation for 1 hr at 37°C. Cells were recovered and washed twice in PBS before lysis in RIPA buffer (50 mM Tris-HCl pH 7.5, 150 mM NaCl, 1.0% NP-40, 0.5% sodium deoxycholate, 0.1% SDS, 1x protease and phosphatase inhibitor (Cell Signaling), and 250 units/ml benzonase nuclease (Sigma)). 40 μg of total proteins were loaded on a 10%–20% SDS-PAGE (Life Technologies) and transferred to a nitrocellulose membrane, 0.45 µm (BioRad). Total protein in each lane was assessed by ponceau S (ThermoFisher) staining and recorded before autoradiography. In all experiments, a replicate of the control lane was treated with chloramphenicol (50 μg/mL) to ensure the mitochondrial origin of the 35S signal. The name associated with each band is proposed based on their relative abundance and molecular weight and is informed by prior studies (Fernández-Silva et al., 2007).

### Genetic interaction screening

#### Screening

K562 cells were infected with pRDA_186 lentiviral vectors, which express sgRNA against a control locus or FXN, blasticidin resistance from the PGK promoter, and a 2A site-expressing EGFP. Cells were sorted for low GFP expression and expanded.

For the screen, the all-in-one Brunello barcoded library was utilized. This library contains 77,441 sgRNA; an average of 4 guides per gene and 1000 non–targeting control guides. Infections were performed in distinct duplicate at a predetermined MOI of ∼0.5 in 12-well plates with 5µg/mL polybrene supplementation. Cells were infected for 2h under centrifugation at 1000 x g, incubated for 22h under standard culturing conditions and pooled 24 h post-centrifugation. Infections were performed with 6.5×10^7^ cells per replicate, in order to achieve a representation of at least 300 cells per sgRNA following puromycin selection. 48 hours after infection, cells were selected with puromycin for 2 days to remove uninfected cells. Cells were passaged in fresh media every 2–3 days. Cells were harvested 11 days after initiation of treatment.

For all screens, genomic DNA (gDNA) was isolated using the XL Maxi NucleoSpin Blood kit (Macherey-Nagel) according to the manufacturer’s protocol. PCR and sequencing were performed as previously described (Doench et al., 2016; Piccioni et al., 2018). Samples were sequenced on a MiSeq (Illumina).

#### Analysis

Analysis of genetic interactors was performed as previously described (To et al. 2019). Briefly, raw sgRNA read counts were normalized to reads per million and then log2 transformed using the following formula:

Log2(Reads from an individual sgRNA/Total reads in the sample x 10^6^ + 1)

Log2 fold-change of each sgRNA was determined relative to guide abundance in the lentiviral library. The replicates were paired throughout the course of the screens.

For each gene and for each replicate, the mean log2 fold-change in the abundance of all 4 sgRNAs was calculated. Log2 fold-changes were averaged by taking the mean across replicates. For each treatment, a null distribution was defined by the 3,726 genes with lowest expression (log2 FPKM = −7) according to publicly available K562 RNA-seq dataset (sample GSM854403 in GEO series GSE34740). To score each gene, its mean log2 fold-change across replicates was Z-score transformed, using the statistics of the null distribution defined as above.

### Polyacrylamide gel electrophoresis and protein immunoblotting

2-5×10^6^ cells were harvested, washed in cold PBS and lysed for 10min on ice in RIPA lysis buffer (50 mM Tris-HCl pH 7.5, 150 mM NaCl, 1.0% NP-40, 0.5% sodium deoxycholate, 0.1% SDS, 1x HALT protease phosphatase inhibitor (Thermo Fisher Scientific), and Pierce Universal Nuclease for Cell Lysis (Thermo Fisher Scientific). Lysates were further clarified by centrifugation for 10min at 16000 x g at 4C. Protein concentration was measured using Pierce 660nm Protein Assay (Thermo Fisher Scientific). 30ug was loaded per well. Electrophoresis was carried out on Novex Tris-Glycine 4-20% or 10-20% gels (Life Technologies) before transfer on a nitrocellulose membrane, 0.45 µm (BioRad). Membranes were blocked for 60mins with Odyssey Blocking Buffer (LI-COR Biosciences) at RT. Membranes were then incubated with primary antibody, diluted in 3%BSA, for 1h at RT or overnight at 4C. Membranes were then washed at RT 5 times in TBST for 5 mins. The membrane was incubated with goat α-rabbit or α-mouse conjugated to IRDye800 or to IRDye680 (LI-COR Biosciences), diluted in 5% milk, for 1h at RT. Membranes were washed 3 times in TBST for 5mins and were scanned for infrared signal using the Odyssey Imaging System (LI-COR Biosciences). Band intensities were analyzed with Image Studio Lite (LI-COR Biosciences).

### Antibodies

Antibodies used here were:

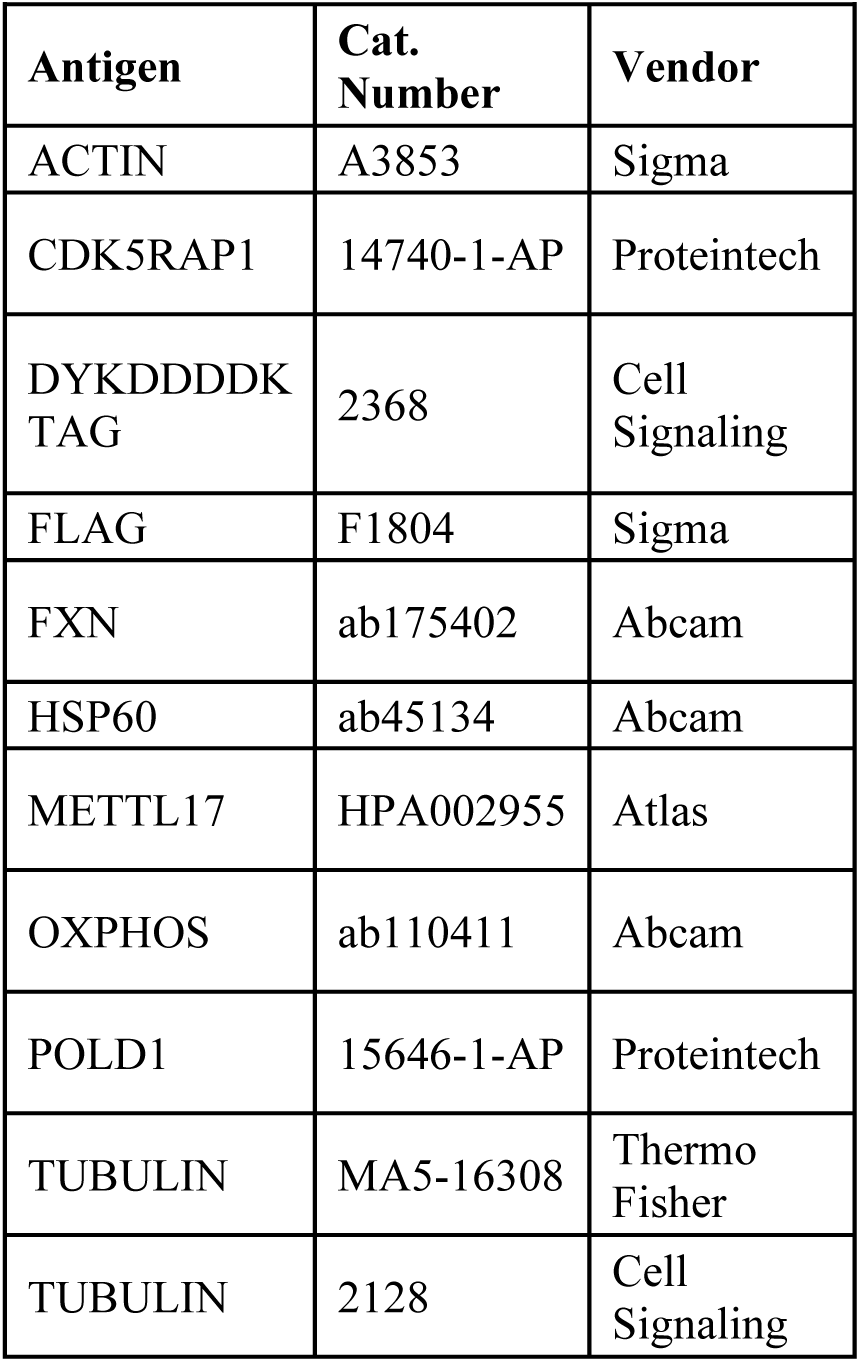

| **Antigen** | **Cat. Number** | **Vendor** |
| --- | --- | --- |
| ACTIN | A3853 | Sigma |
| CDK5RAP1 | 14740-1-AP | Proteintech |
| DYKDDDDK TAG | 2368 | Cell Signaling |
| FLAG | F1804 | Sigma |
| FXN | ab175402 | Abcam |
| HSP60 | ab45134 | Abcam |
| METTL17 | HPA002955 | Atlas |
| OXPHOS | ab110411 | Abcam |
| POLD1 | 15646-1-AP | Proteintech |
| TUBULIN | MA5-16308 | Thermo Fisher |
| TUBULIN | 2128 | Cell Signaling |

### Cell viability assay in galactose vs. glucose

To measure viability in galactose vs glucose, cells were washed in PBS, counted and an equal number of cells was seeded in culture media containing 25mM glucose or 25mM galactose with 10% dialyzed FBS. 24h later, cells were collected and viable cells were determined using a Vi-Cell Counter (Beckman).

### qPCR

2.5×10^6^ cells were collected per sample. RNA was extracted from total cells with an RNeasy plus kit (QIAGEN) and DNase-I digested before murine leukemia virus (MLV) reverse transcription using random primers (Promega). qPCR was performed using the TaqMan technology (Life Technologies), using probes Hs02596859_g1 (12S), Hs02596860_s1 (16S), Hs00224159_m1 (METTL17) and Hs00427620_m1 (TBP). All data were normalized to TBP.

### Formaldehyde RNA immunoprecipitation (fRIP) assay to METTL17-12S binding

fRIP assay was performed as previously described (Hendrickson et al, 2016) with some minor changes. Briefly, 5×10^6^ K562 cells were cross-linked with 0.1% formaldehyde (Polysciences, 04018) and slowly rotated for 10 min at room temperature. To quench this reaction, glycine was added to a final concentration of 125mM and incubated for 5 min at room temperature with rotation. Cells were collected and spun down at 300g for 5min at 4°C. The supernatant was removed and the pellet was washed with 1mL cold PBS. The supernatant was removed and the pellet was resuspended in 1ml RIPA buffer (buffer (50 mM Tris-HCl pH 7.5, 150 mM NaCl, 1.0% NP-40, 0.5% sodium deoxycholate, 0.1% SDS, 1x protease and phosphatase inhibitor (Cell Signaling), and 250 units/ml benzonase nuclease (Sigma)). The cells were sonicated for 3 times each (10 sec per cycle, amplitude 7 μm) with a sonifier (Branson). Samples were then further lysed on ice for 10 min. To remove cell debris, the extract was centrifuged at 12,000g for 10 min at 4°C. 150 μL of the supernatant was removed and stored at −20°C (input sample), and the remainder of the sample was used for precipitation. Anti-FLAG® M2 magnetic beads (Sigma) were equilibrated 3 times with 1 mL RIPA buffer. 20μL of anti-FLAG beads were used for precipitation. The IP was performed overnight at 4°C on a spinning wheel.

Beads were recovered after extensive washing in RIPA buffer. After the last washing step, the supernatant was removed, 100μL of RNAse free water with RNAsin (40 U/mL) was added and samples incubated in a thermomixer (Eppendorf) at 300 rpm for at 55°C for 1h for decrosslinking. 1ml of Trizol (Invitrogen) was then added to each sample, and RNA extraction was performed per the manufacturer’s recommendations. RNA samples were subsequently quantified for 12S levels by qPCR, and the values were normalized to input 12S and FLAG tagged protein levels.

### mtDNA copy number

1×10^5^ cells from each condition (n = 3) were harvested and lysed in 100μL mtDNA lysis buffer (25mM NaOH, 0.2mM EDTA) before incubation at 95°C for 15min. 100μL of 40mM Tris-HCl pH 7.5 was added to neutralize the reaction on ice. Samples were diluted 50x and the ratio between mitochondrial and nuclear DNA was determined using a custom Taqman based assay and qPCR using a CFX Opus 384 quantitative PCR machine (Biorad). Relative mtDNA abundance was determine using the ΔΔCt method.

### Mitostring

2×10^6^ cells were collected per sample. RNA was extracted from total cells with an RNeasy kit (QIAGEN). MitoString is a mitochondrial-specific version of NanoString and was performed as previously described (Wolf and Mootha, 2014). All counts were normalized to TUBB.

### Mitochondrial isolation

5×10^7^ cells were harvested, washed in PBS, and washed once with 10 mL of MB buffer (210 mM mannitol, 70 mM sucrose, 10 mM HEPES-KOH at pH 7.4, 1mM EDTA, protease/phosphatase inhibitor). Cells were resuspended in 1 mL of MB buffer supplemented with 1x HALT protease phosphatase inhibitor (Thermo Fisher Scientific) and transferred to 2 mL glass homogenizer (Kontes). Cells were broken with ∼35 strokes of a large pestle on ice. Sample volume was then increased to 6 mL with MB buffer. The mixture was centrifuged at 2,000xg for 5 min and the pellet (nuclei and intact cells) was discarded. The supernatant was centrifuged at 13,000xg for 10 min at 4°C. The mitochondrial pellets were washed with MB buffer once, and resuspended in RIPA lysis buffer.

### Oxygen Consumption

1.50×10^5^ K562 cells per well were plated on a Seahorse plate coated with Cell-Tak Cell and Tissue Adhesive (Corning Life Sciences) in XF DMEM (Agilent) containing 5.55mM glucose, 4mM Glutamine, 1mM sodium pyruvate, and oxygen consumption was recorded using a Seahorse XF96 Analyzer (Seahorse Biosciences). Each measurement was performed over 6 min after a 3 min mix and a 3 min wait period. Basal measurements were collected 4 times, followed by 4 measurements after addition of oligomycin (final concentration 2µM), followed by 4 measurements after addition of Bam15 (final concentration 3µM), followed by 4 measurements after addition of Piericidin A+ Antimycin A (final concentration 0.5 µM).

### Proliferation assays

Cell proliferation assays were performed 7-8 days following lentiviral infection. Cells were seeded at an initial density of 1×10^5^ cells/mL and cultured for 3 days. Viable cell number was then determined using a Vi-Cell Counter (Beckman).

### Expression and purification of METTL17-8XHIS

A human METTL17 construct was designed for expression in E. coli in which the N-terminal mitochondrial targeting sequence was removed and a C-terminal TEV site and 8X HIS-tag were introduced. The METTL17-8XHIS gene was codon optimized for *E. coli* and ligated into a pET-30a(+) expression vector. The protocol for overexpression and purification of METTL17-8XHIS was adapted from that used for the yeast homolog (Alam et al., 2021). METTL17-8XHIS was overexpressed in OverExpress C41(DE3) cells in LB supplemented with 50 μM FeCl3 and 300 μM L-cysteine at 37°C. Cells were grown until an OD600 of 0.8 and then expression was induced with 1mM IPTG. Cells were pelleted after 4 hours, then frozen and stored in liquid nitrogen until purification. At the time of purification, pellets were thawed and resuspended in Buffer A containing 500 mM NaCl, 5 mM imidazole, 5% glycerol, 1 mM TCEP, 40 mM HEPES pH 7.4, and supplemented with lysonase, benzonase, DNase, ribonuclease A, Roche Complete protease, and PMSF. The cells were lysed using sonication and clarified through centrifugation at 40,000 × g for one hour. METTL17-8XHIS was then purified over TALON cobalt resin. After washing the column with Buffer A containing 40 mM imidazole, the protein was washed with 30 CVs of a solution containing 2.5 M NaCl, 5% glycerol, 1 mM TCEP, 40 mM HEPES pH 7.4, and 0.1 mg/mL ribonuclease A and DNase I to remove nucleic acid contamination. The protein was eluted with Buffer A containing 300 mM imidazole and then run over a PD10 column into a buffer containing 500 mM NaCl, 5% glycerol, 1 mM TCEP, and 40 mM HEPES pH 7.4. Sizing of the protein was done over a Superdex 200 increase 10/300 GL column. The variant CYS^Mut^-8xHIS containing the mutations C333S, C339S, C347S, and C404S was expressed and purified using the same method.

### UV/vis spectroscopy

UV/vis spectra were collected at room temperature with a Cary 50 UV/vis spectrophotometer using a 1 cm pathlength quartz cuvette. Aerobically purified METTL17-8XHIS or CYS^Mut^-8xHIS was transferred into a Coy Labs glovebox (<5 ppm O_2_). The protein solution was equilibrated to the glovebox atmosphere for at least 30 min on ice before it was centrifuged for 3 min at 14,000 × g and room temperature, then buffer-exchanged into an anaerobic buffer containing 25 mM HEPES, pH 7.5, 10% glycerol, 500 mM NaCl using a PD-10 column (GE Healthcare). Samples were diluted (if necessary) with the same buffer and transferred to a quartz cuvette.

### EPR spectroscopy

Electron paramagnetic resonance (EPR) spectra were recorded on a Bruker EMX spectrometer at 9.37 GHz as frozen solutions. Samples were transferred to a quartz tube (made from clear fused quartz tubing of 3 mm I.D. & 4 mm O.D. from Wale Apparatus) in a Coy Labs glovebox (<5 ppm O_2_), capped with a rubber septum, and frozen in liquid N_2_ outside the glovebox. Each sample contained approximately 200 μL of 30 μM aerobically purified METTL17-8XHIS with/without 2 mM Na_2_S_2_O_4_ as a reductant or 0.5 mM indigodisulfonic acid as an oxidant.

### Fe analysis

Protein concentrations were determined by BCA assay using bovine serum albumin (Thermo Scientific) as standards (Smith et al., 1985). Fe concentrations were determined by inductively coupled plasma mass spectrometry (ICP-MS). ICP-MS data were recorded on an Agilent 7900 ICP-MS instrument. Samples were prepared by first digesting the protein in 70% nitric acid (TraceMetalTM Grade, Fischer) at 60 °C, and then diluting it with Milli-Q water such that the final concentration of nitric acid is 2%. Standards for Fe were prepared from a 100 ppm transition metal standard solution (Specpure, Thermo Fischer). All samples and standards contained 1 ppb Tb (final concentration) as an internal standard.

### Mitoribosome purification for structural studies

Yeast mitochondria were harvested as previously described (Amunts *et al*., 2014). Four litres of *S. cerevisiae* were grown in YPG media (1% yeast extract, 2% peptone, 3% glycerol) until an optical density at 600 nm (OD_600_) of 1.8. The cells were then centrifuged at 4,500 × *g,* for 9 min, the pellet washed with pre-cooled distilled water and further centrifuged for 15 min at 4,500 × *g.* The pellet was resuspended in pre-warmed dithiothreitol (DTT) buffer (100 mM Tris-HCl pH 9.3, 10 mM DTT). The cells were pelleted by centrifugation at 3,500 × *g,* for 10 min and resuspended in Zymolyase buffer (20 mM K_2_HPO_4_-HCl pH 7.4, 1.2 M sorbitol). 1 mg ZymolyaseF100T (MP Biomedicals, LLC) was added per gram wet weight and the solution was shaken slowly at 30°C for 60 min. This was followed by centrifugation at 4,000 × *g,* for 15 min. The pellet was further resuspended in Zymolyase buffer and centrifuged for a further 15 min at 4,000 × *g.* The pellet was then resuspended in homogenization buffer (20 mM HEPES-KOH pH 7.45, 0.6 M sorbitol, 1 mM EDTA) and lysed with 15 strokes in a glass homogenizer. To separate the cell debris and nuclei from mitochondria the solution was centrifuged at 2000 × *g,* for 20 min and the supernatant collected, followed by further centrifugation at 4500 × *g,* for 20 min. Crude mitochondria were pelleted at centrifugation at of 13,000 × *g,* for 25 min, and further purified on a sucrose gradient in SEM buffer (250 mM sucrose, 20 mM HEPES-KOH pH 7.5, 1 mM EDTA) by ultracentrifugation at 141,000 x *g* for 1 hour. Mitochondrial samples were pooled and lysed in a buffer of 25 mM HEPES-KOH pH 7.5, 100 mM KCl, 25 mM Mg(OAc)_2_, 1.7% Triton X-100, 2 mM DTT). Upon centrifugation at 30,000 × *g,* for 20 min, the supernatant was loaded on a 1 M sucrose cushion in a buffer containing 20 mM HEPES-KOH pH 7.5, 100 mM KCl, 20 mM Mg(OAc)_2_, 1% Triton X-100, 2 mM DTT. The pellet was resuspended in 10 mM Tris-HCl, pH 7.0, 60 mM KCl, 60 mM NH_4_Cl, 10 mM Mg(OAc)_2_, and applied on a 15-40% sucrose gradient prepared in the same buffer. The fractions containing the SSU were collected, sucrose removed and buffer exchanged to 20 mM HEPES-KOH pH 7.5, 50 mM KCl, 5.0 mM Mg(OAc)_2_, 2.0 mM DTT and 0.05% n-dodecyl-β-D-maltoside by passing through a concentrator (Vivaspin) with a 30 kDa molecular weight cutoff. The final concentration of the mitoribosomes measured at an optical density at 280 nm (OD280) was 3.6

### Production of mtIF3

The sequence of mtIF3/AIM23 gene of yeast *S. cerevisiae* was inserted in pET30a vector between Nde I and Xho I restriction sites. The plasmid was used to transform *E. coli* Rosetta strain. The culture was inoculated by diluting 100 x with LB and grown until OD600 reached the value of 0.8. Protein production was induced by 0.25 mM IPTG at 30 °C for 3 hours. Biomass was collected by centrifugation at 3500x *g* for 5 min and resuspended in native lysis buffer (25 mM NaHPO4, 500 mM NaCl, 25 mM imidazol, pH 7.4). The suspension was sonicated 6 x 10 sec with 20 sec breaks inbetween. The debris pelleted by centrifugation at 30000x *g* for 20 min. The mtIF3 was then purified on HisTrap HP column by metallo-chelating chromatography. The second step of purification and buffer exchange was done on Superdex 75 10/300 GL column in 20 mM HEPES-KOH, 200 mM NaCl, pH 7.0. The fractions were analysed by SDS-PAGE, pooled and concentrated on Vivaspin20 concentrators to 85 ∝M. Purified mtIF3 was added to the SSU in 5x molar excess and incubated for 30 min.

### Cryo-EM data collection and image processing

For cryo-EM, 3 μL of ∼100 nM (A_260_ 4.0 or 6.0) mitoribosome sample was applied onto a glow-discharged (20 mA for 30 sec) holey-carbon grid (Quantifoil R2/1 copper, mesh 300) coated with continuous carbon (of ∼3 nm thickness) and incubated for 30 sec in a controlled environment of 100% humidity and 4°C temperature. The grids were blotted for 3.5 sec, followed by plunge-freezing in liquid ethane, using a Vitrobot MKIV (FEI/Thermofischer). The datasets were collected on FEI Titan Krios (FEI/Thermofischer) transmission electron microscope operated at 300 keV with a slit width of 20 eV on a GIF quantum energy filter (Gatan). A K2 Summit detector (Gatan) was used at a pixel size of 0.83 Å (magnification of 165,000x) with a dose of 32 electrons/Å^2^ fractionated over 20 frames. A defocus range of 1.2 to 2.8 μm was applied.

Beam-induced motion correction was performed for all data sets using RELION 3.0 (Zivanov et al., 2018). Motion-corrected micrographs were used for contrast transfer function (CTF) estimation with GCTF (Zhang, 2016). Particles were picked by Gautomatch (https://www.mrc-lmb.cam.ac.uk/kzhang) with reference-free, followed by reference-aided particle picking procedures. Reference-free 2D classification was carried out to sort useful particles from falsely picked objects, which were then subjected to 3D refinement, followed by a 3D classification with local-angular search. UCFS Chimera (Pettersen et al., 2004) was used to visualize and interpret the maps. 3D classes corresponding to unaligned or low-quality particles were removed. Well-resolved classes were pooled and subjected to 3D refinement and CTF refinement (beam-tilt, per-particle defocus, per-micrograph astigmatism) by RELION 3.1 (Zivanov et al., 2020), followed by Bayesian polishing. Particles were separated into multi optics groups based on acquisition areas and date of data collection. Second round of 3D refinement and CTF refinement (beam-tilt, trefoil and fourth-order aberrations, magnification anisotropy, per-particle defocus, per-micrograph astigmatism) were performed, followed by 3D refinement.

To classify the factor binding states, a non-align focus 3D classifications with particle-signal subtraction using the mask covering the factor binding were done with RELION 3.1. The particles of each state were pooled, subtracted signal was reverted and 3D refinement was done with the corresponding solvent mask. To improve the local resolution, the several local masks were prepared and used for local-masked 3D refinements. Nominal resolutions are based on gold-standard, applying the 0.143 criterion on the Fourier Shell Correlation (FSC) between reconstructed half-maps. Maps were subjected to B-factor sharpening and local-resolution filtering by RELION 3.1, superposed to the overall map and combined for the model refinement.

### Model building, refinement, and analysis

The starting models for SSU was PDB ID 5MRC (Desai *et al*., 2017). These SSU model was rigid body fitted into the maps, followed by manual revision. Initial model of mtIF3 was generated by SWISS-MODEL (Waterhouse et al., 2018), while that of METTL17 was built manually. Ligands and specific extensions/insertions were built manually based on the density. *Coot* 0.9 with Ramachandran and torsion restraints (Emsley et al., 2010) was used for manual fitting and revision of the model. Geometrical restraints of modified residues and ligands were calculated by Grade Web Server (http://grade.globalphasing.org). Final models were subjected to refinement of energy minimization and ADP estimation by Phenix.real_space_refine (Liebschner et al., 2019) with rotamer restraints without Ramachandran restrains, against the composed maps with B-factor sharpening and local-resolution filtering. Reference restrains was also applied for non-modified protein residues, using the input models from *Coot* as the reference. Metal­ coordination restrains were generated by ReadySet in the PHENIX suite and used for the refinement with some modifications. Refined models were validated with MolProbity (Williams et al., 2018) and EMRinger (Barad et al., 2015) in the PHENIX suite. UCSF ChimeraX 0.91 (Goddard et al., 2018) was used to make figures.

## Supplemental Figures and Legends

**Fig S1:**
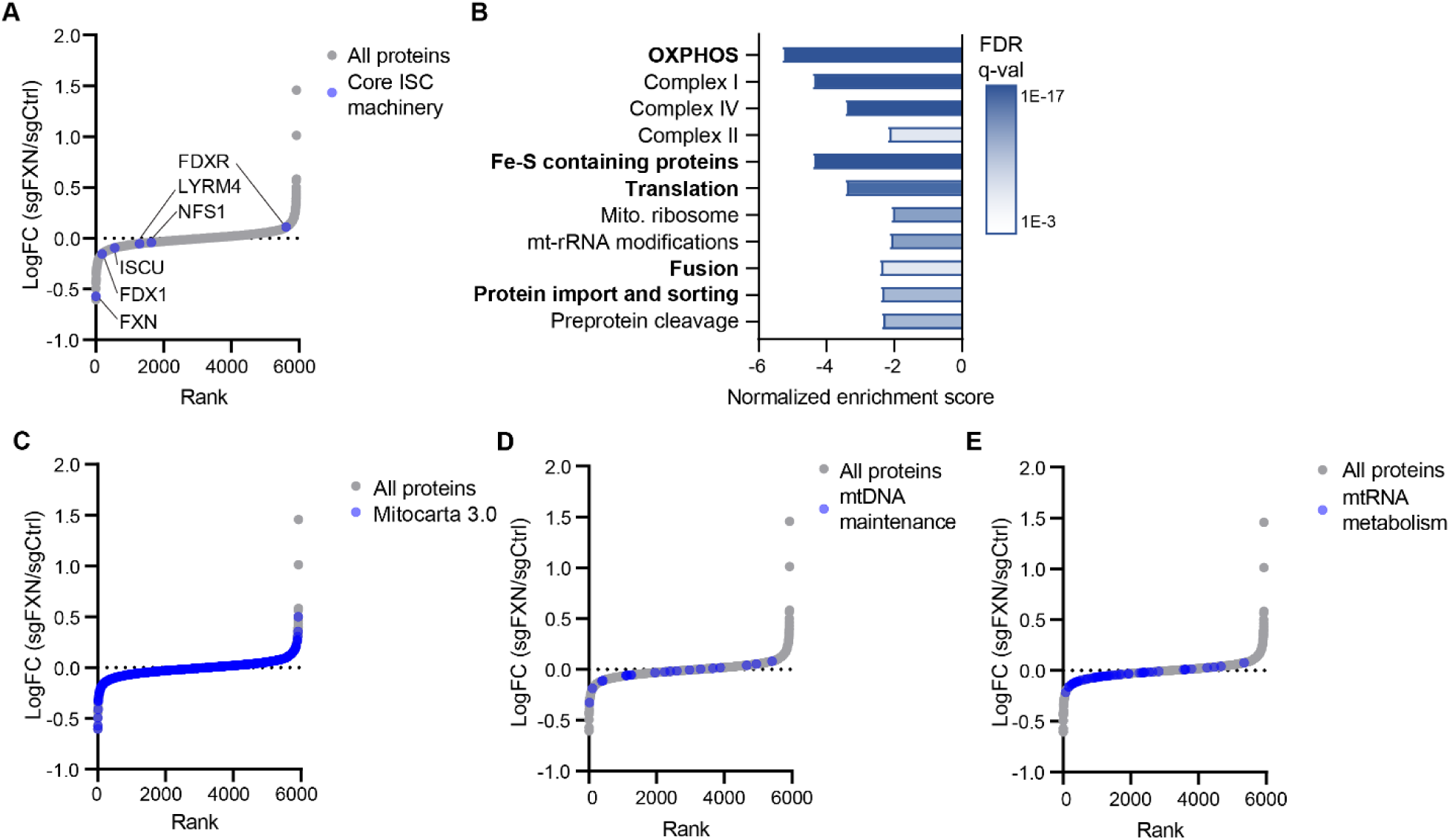
OXPHOS, but not ISC machinery or the mito-proteome, is depleted in the absence of FXN. A. Waterfall plots of protein fold change in FXN/Control cells, highlighting proteins in the core ISC machinery. B. Analysis of MitoPathways depleted in FXN null cells. Pathways in bold are independent. C-E. Waterfall plots of protein fold change in FXN/Control cells, highlighting proteins in MitoCarta 3.0 (B) mtDNA maintenance proteins (D) and mtRNA metabolism (E).

**Fig S2:**
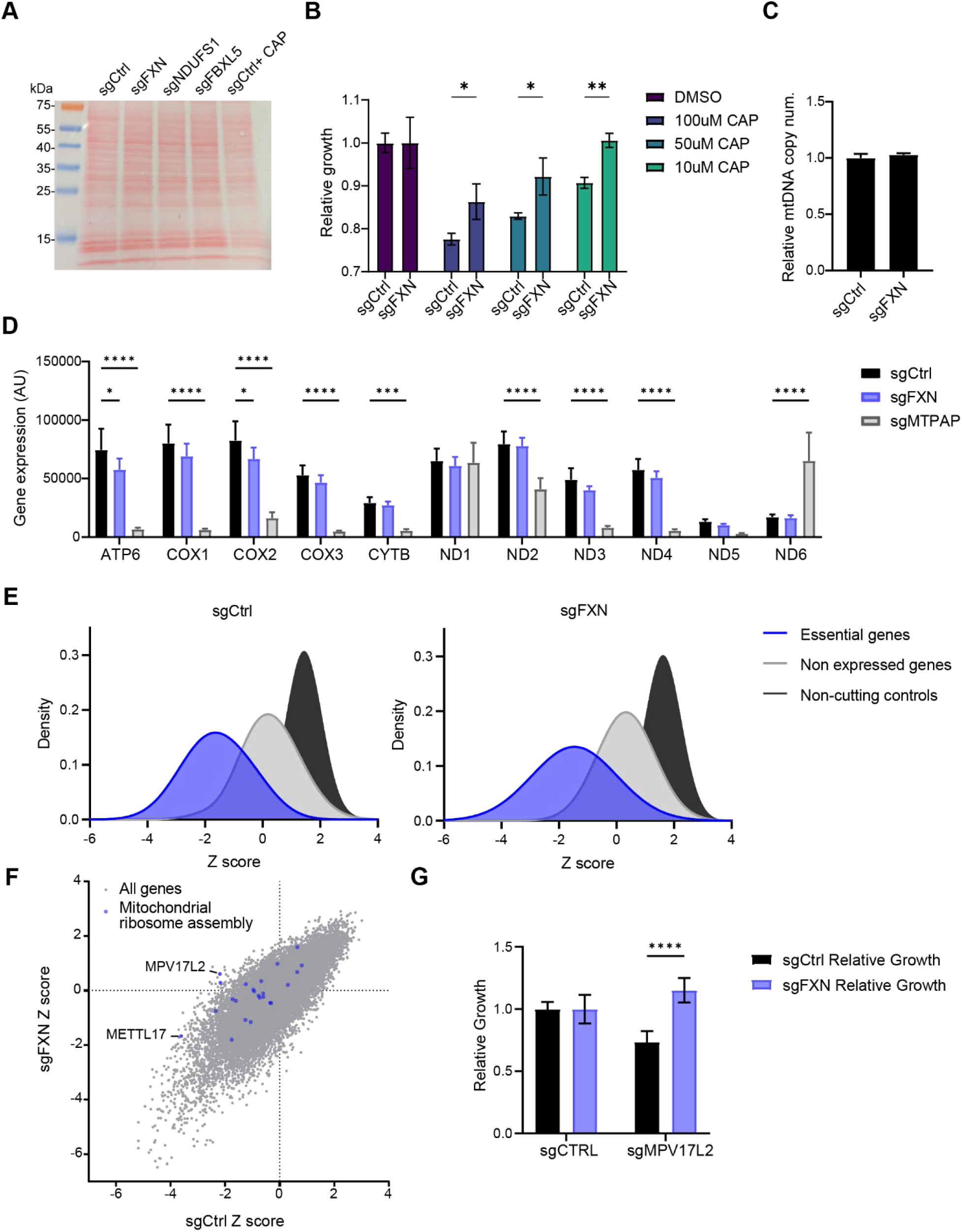
mtDNA replication and transcription is not significantly altered in FXN null cells. A. Ponceau S staining of the protein membrane found in Fig 2A. B. sgCtrl and sgFXN cells were grown for 72h in escalating concentrations on chloramphenicol, and the relative growth of each strain compared to DMSO treatment was calculated. C. qPCR for mtDNA copy number in sgCtrl and sgFXN cells. D. Mitostring assay examining the levels of mtDNA encoded transcripts in sgCtrl, sgFXN and sgMTPAP cells. E. Histograms of the Z score of cutting controls, non-expressed genes and essential genes as defined by (Hart et al., 2015) in the genetic interaction screens preformed in sgCtrl or sgFXN cells. F. Scatterplot of Z scores showing knockouts growth in sgCtrl vs. sgFXN backgrounds. All the mitochondrial ribosome assembly genes, as defined by MitoCarta 3.0, are highlighted in blue. G. Relative growth rates of cells edited for control or MPV17L2 gene, on the background on sgCtrl or sgFXN. Growth rates were normalized to the unedited growth rate for each strain. All bar plots show mean ± SD. *=p < 0.05, **=p < 0.01, ***=p < 0.001, ****=p < 0.0001. One-way ANOVA with Bonferroni’s post-test.

**Fig S3:**
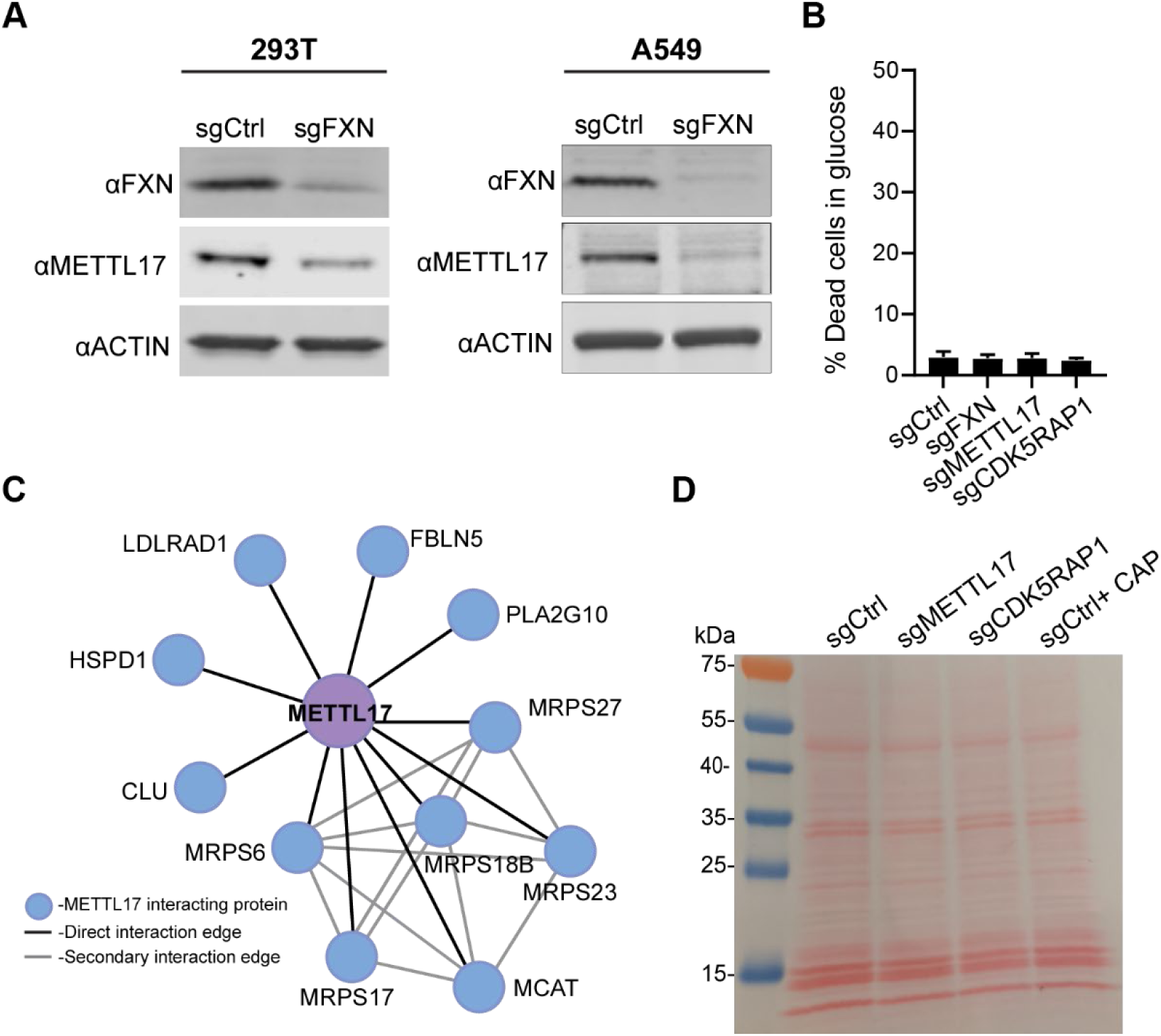
METTL17 is depleted in FXN null cells and is linked to mitochondrial translation. A. Immunoblot for FXN, METTL17 and the loading control actin in 293T and A549 cells edited with control or FXN guides. B. Cells edited for control, FXN, METTL17 and CDK5RAP1 genes were grown for 24h in glucose media, and viability was assessed on each background. C. Protein-protein interactions identified for METTL17 in 293T cells as identified by (Huttlin et al, 2021). D. Ponceau S staining of the protein membrane found in Fig 3E. All bar plots show mean ± SD.

**Fig S4:**
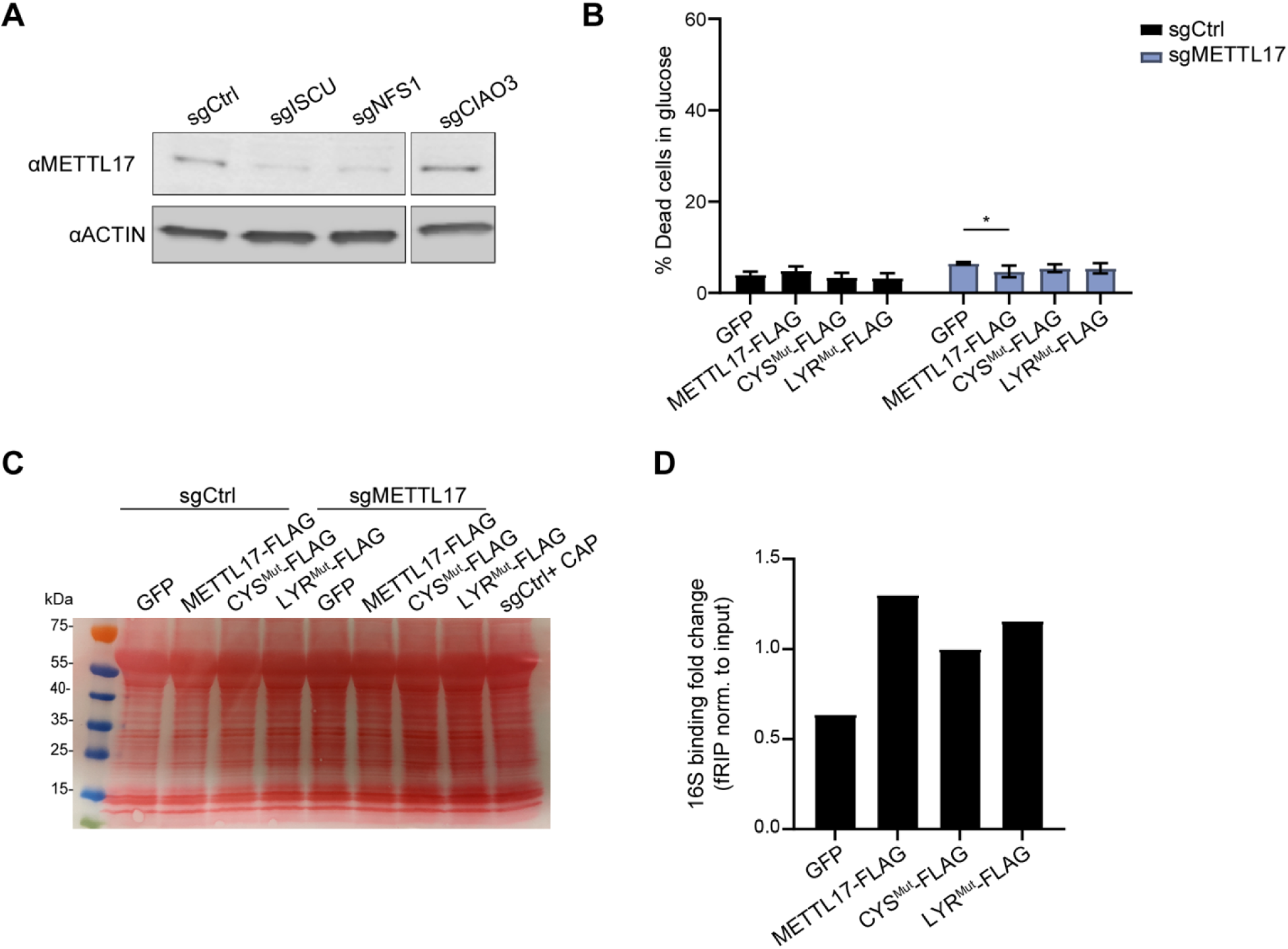
METTL17 has characteristics of an Fe-S cluster binding protein. A. Immunoblot for METTL17 and the loading control actin in cells edited for control, ISC genes (ISCU and NFS1) or a CIA gene (CIAO3). B. Cells edited for control or METTL17 genes and expressing WT or mutant forms of METTL17 were grown for 24h in glucose media, and viability was assessed on each background. C. Ponceau S staining of the protein membrane found in Fig 4E. D. Formaldehyde-linked RNA immunoprecipitation of the 16S to GFP, METTL17-FLAG, CYSMut-FLAG or LYRMut-FLAG proteins. Results were normalized to input construct and 16S levels. All bar plots show mean ± SD. *=p < 0.05, One-way ANOVA with Bonferroni’s post-test.

**Fig S5:**
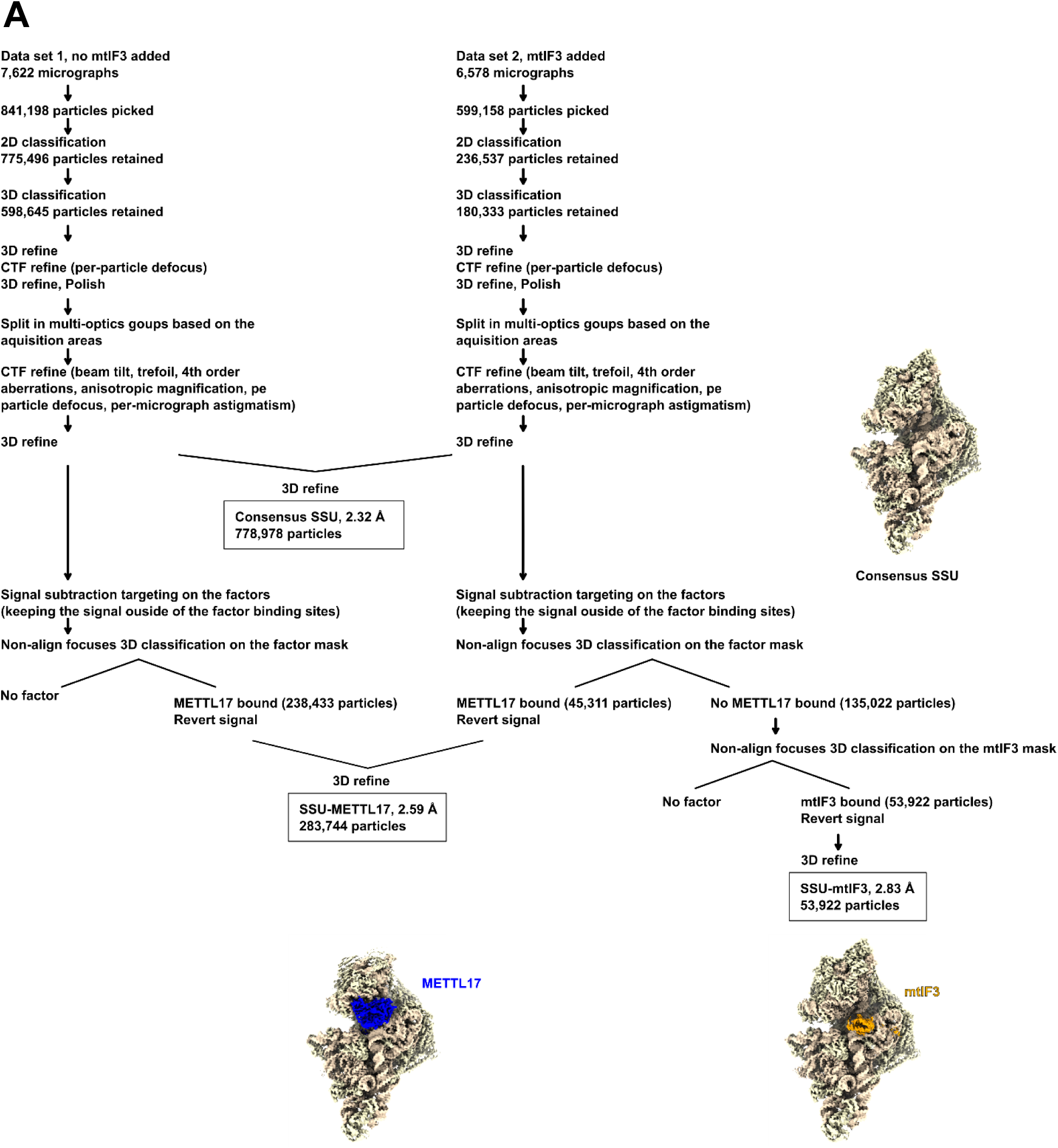
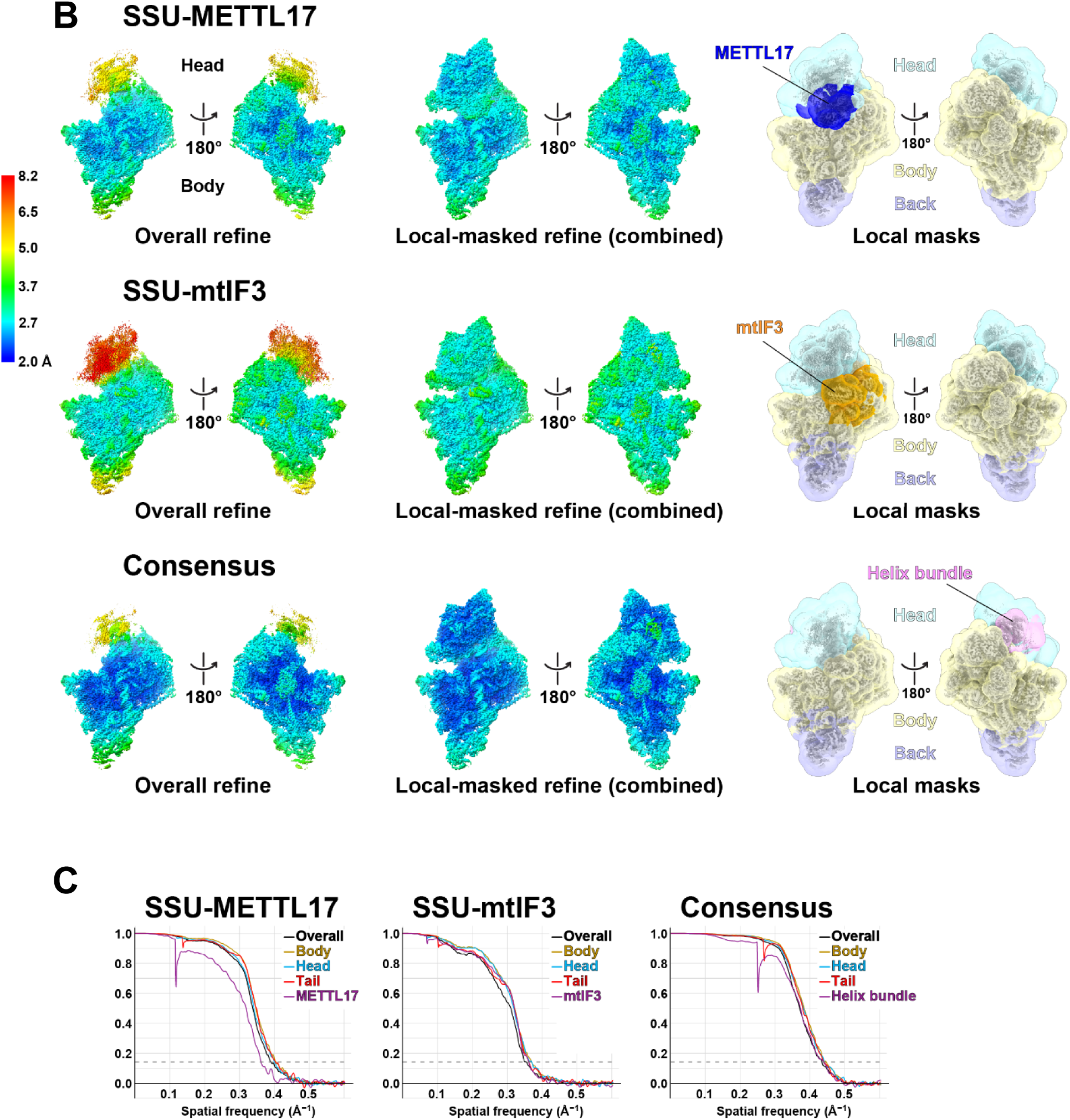
Cryo-EM image processing. A. The data processing scheme. B. Overall maps, combined maps of the local-masked refinements colored by local resolution are shown for SSU-I:v1ET1L17 (top), SSU-mtiF3 (middle), and SSU (bottom). C. Fourier Shell Correlation curves of the half maps and local-masked refinements. The 0.143 criterion is shown as dashed lines.

**Fig S6:**
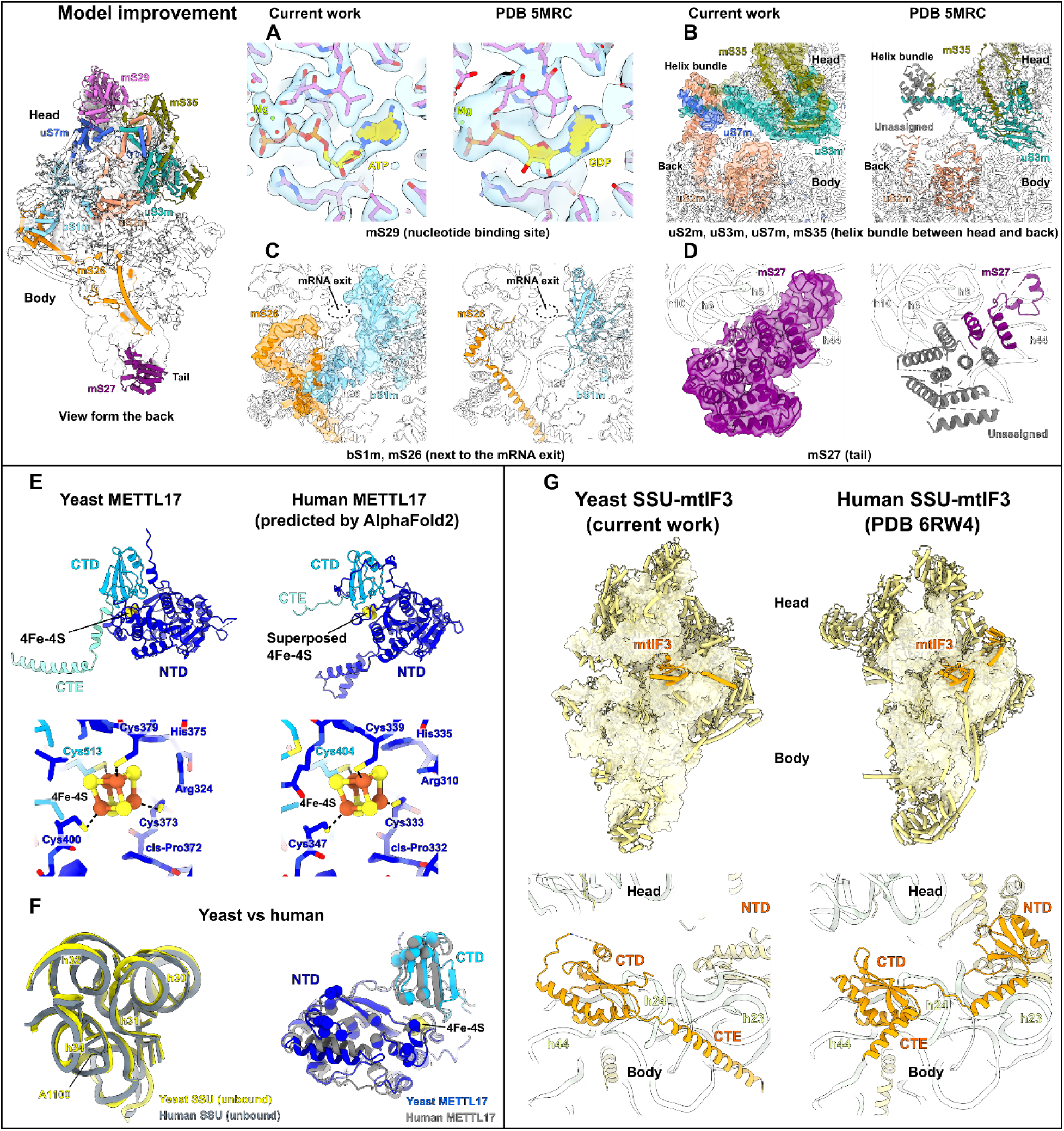
Cryo-EM structure of the yeast SSU, METTL17 and mtIF3. A-D. Improvements in the model of *S. cerevisiae* mitoribosome. Overview of the SSU model from the back with improved proteins colored. The close-up views show modeled elements with their corresponding density map, and equivalent regions from previous studies (Desai *et al*., 2017) are shown for comparison. A. The nucleotide density for mS29 in the SSU head. B. The density and corresponding models of uS2m, uS3m, uS7m, mS35 that form a previously unsigned helix bundle between the head and body. C. Complete models for bS1m and mS26 that form contacts at the mRNA channel exit. D. Remodeled and reannotated mS27 interacts with h44, which was previously partially built as poly-Ala and named mS44. E. Comparison between the yeast cryo-EM model and human *AlphaFold2* (Jumper *et al*., 2021) prediction of METTL17 shows that the predicted conformations of the NTD (blue) and CTD (light blue) are highly similar, including the coordination of the 4Fe-4S shown in the close-up view, and structural differences are observed only in the terminal extensions. The Fe-S cluster in the human model was placed by superposing that of the yeast cryo-EM structure. F. Comparison between yeast and human SSU (left) and METTL17 (right) interfaces. Phylum-specific protein extensions have been removed for clarity. The residues involved in interactions are shown in sticks for RNA and spheres for protein. G. Comparison between yeast and human (Khawaja *et al*., 2020) SSU-IF3 complex with close-up views showing that the binding of the mtIF3 CTD (orange) is conserved. Thus, human mtIF3 has similar structural characteristics and would also clash with METTL17 on the SSU. The NTD of mtIF3 is not well resolved in our map, and thus hasn’t been modelled. On the other hand, the C-terminal extensions (CTE) forming a helix have different orientations. The CTE in yeast keeps contacting the rRNA in the body, whereas that of human is exposed.

**Fig S7:**
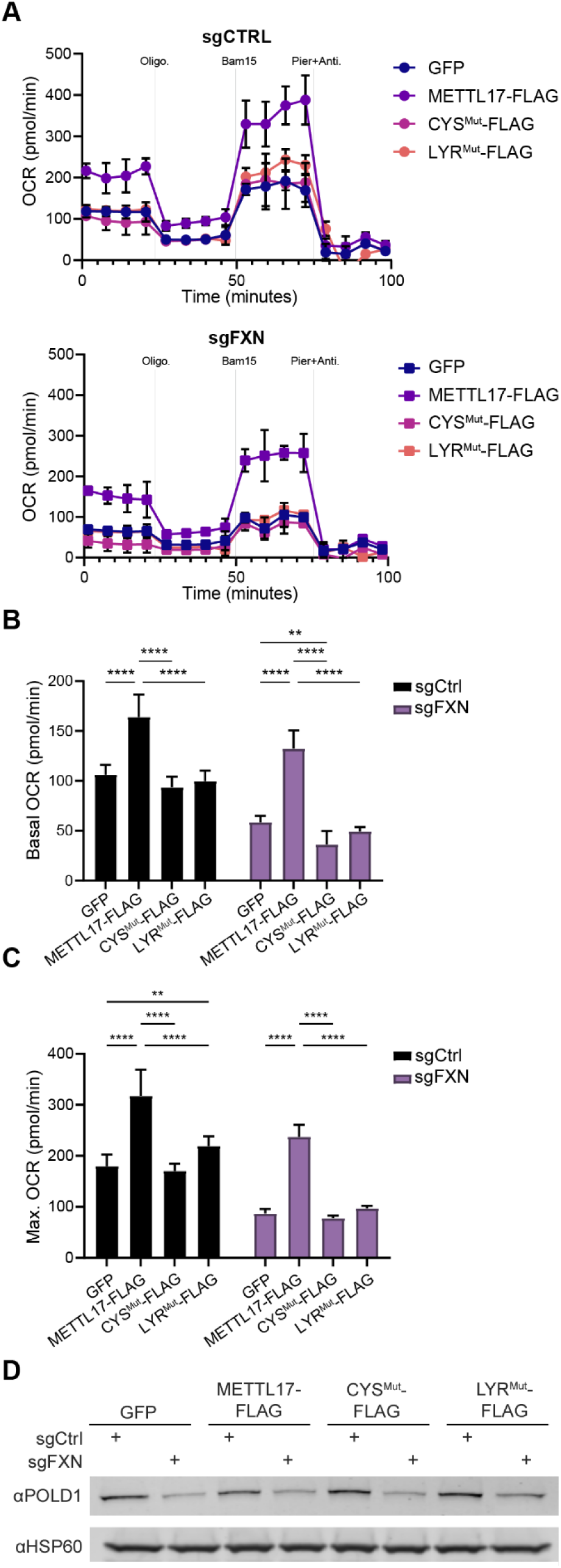
METTL17 overexpression restores the faulty mitochondrial bioenergetics of FXN depleted cells. A. Oxygen consumption rate of Control (top) or FXN (bottom) edited cells expressing GFP, METTL17-FLAG, CYSMut-FLAG or LYRMut-FLAG. Cells were sequentially treated with oligomycin, Bam15 and Piericidin A+ Antimycin A. B-C. Basal (B) and maximal (C) OCR of Control or FXN edited cells expressing GFP or METTL17-FLAG. D. Immunoblots examining POLD1 or the loading control HSP60 in Control or FXN edited cells expressing GFP, METTL17-FLAG, CYS^Mut^-FLAG or LYR^Mut^-FLAG constructs. All bar plots show mean ± SD. **=p < 0.01, ****=p < 0.0001. One-way ANOVA with Bonferroni’s post-test.

**Table S1.** Data collection, processing, model refinement and validation statistics.

|  | SSU-METTL17 | SSU-mtIF3 | Consensus SSU |
| --- | --- | --- | --- |
| **Data collection and Processing** |  |  |  |
| Microscope | Titan Krios | Titan Krios | Titan Krios |
| Voltage (kV) | 300 | 300 | 300 |
| Camera | K2 Summit | K2 Summit | K2 Summit |
| Magnification | 165,000 | 165,000 | 165,000 |
| Pixel size at detector (Å/pixel) | 0.83 | 0.83 | 0.83 |
| Total electron exposure (e<sup>-</sup>/Å<sup>2</sup>) | 32 | 32 | 32 |
| Exposure rate (e<sup>-</sup>/pixel/sec) | 5.6 | 5.6 | 5.6 |
| Number of frames | 20 | 20 | 20 |
| Defocus range (μm) | -1.2 to -2.8 | -1.2 to -2.8 | -1.2 to -2.8 |
| Energy filter slit width (V) | 20 | 20 | 20 |
| Micrographs collected (no.) | 14,200 | 6,578 | 14,200 |
| Final particles (no.) | 283,744 | 53,922 | 778,978 |
| Point group | C<sub>1</sub> | C<sub>1</sub> | C<sub>1</sub> |
| Resolution (global, Å) FSC 0.143 (masked) | 2.59 | 2.83 | 2.32 |
| Map-sharpening *B* factor (Å<sup>2</sup>) | -41 | -37 | -43 |
| **Model composition** |  |  |  |
| Atoms | 91,232 | 88,476 | 88,581 |
| RNA residues | 1,478 | 1,477 | 1,485 |
| Protein residues (non-modified/ modified) | 7386/ 1 | 7,056/ 1 | 7,038/ 1 |
| Metal ions (Mg<sup>2+</sup>/ K<sup>+</sup>) | 63/ 8 | 58/ 8 | 103/ 21 |
| Ligands (ATP/ 4Fe-4S) | 1/ 1 | 1/ 0 | 1/ 0 |
| Waters | 0 | 3 | 3 |
| **Model Refinement** |  |  |  |
| Model-Map CC (CC<sub>mask</sub>/ CC<sub>box</sub>/ CC<sub>peaks</sub>/ CC<sub>volume</sub>) | 0.73/ 0.79/ 0.71/ 0.75 | 0.80/ 0.85/ 0.78/ 0.82 | 0.80/ 0.84/ 0.77/ 0.81 |
| Resolution (Å) by model-to-map FSC, threshold 0.50 (masked/ unmasked) | 2.10/ 2.12 | 2.40/ 2.40 | 1.70/ 1.70 |
| Average *B* factor (Å<sup>2</sup>) (RNA/ protein) | 27/ 21 | 29/ 35 | 27/ 23 |
| R.m.s. deviations, bond lengths (Å)/ bond angles (°) | 0.009/ 0.799 | 0.002/ 0.505 | 0.002/ 0.508 |
| **Validation** |  |  |  |
| MolProbity score | 1.79 | 1.32 | 1.34 |
| CaBLAM outliers (%) | 1.40 | 1.02 | 1.02 |
| Clash score | 6.82 | 5.82 | 5.48 |
| Rotamer outliers (%) | 3.04 | 0.73 | 1.16 |
| C<sub>β</sub> deviations | 0 | 0 | 0 |
| Ramachandran plot (%) (Favored/ allowed/ disallowed) | 97.80/ 2.20/ 0.00 | 98.59/ 1.39/ 0.01 | 98.55/ 1.45/ 0.00 |

